# FF12MC: A revised AMBER forcefield and new protein simulation protocol

**DOI:** 10.1101/061184

**Authors:** Yuan-Ping Pang

**Affiliations:** Computer-Aided Molecular Design Laboratory, Mayo Clinic, Rochester, MN 55905, USA

**Keywords:** Protein folding, Protein dynamics, Protein simulation, Protein structure refinement, Molecular dynamics simulation, Force field, Chignolin, CLN025, Trp-cage, BPTI

## Abstract

Specialized to simulate proteins in molecular dynamics (MD) simulations with explicit solvation, FF12MC is a combination of a new protein simulation protocol employing uniformly reduced atomic masses by tenfold and a revised AMBER forcefield FF99 with (*i*) shortened CH bonds, (*ii*) removal of torsions involving a nonperipheral *sp^3^* atom, and (*iii*) reduced 1-4 interaction scaling factors of torsions *ϕ* and *ψ* This article reports that in multiple, distinct, independent, unrestricted, unbiased, isobaric-isothermal, and classical MD simulations FF12MC can (*i*) simulate the experimentally observed flipping between left-and right-handed configurations for C14-C38 of BPTI in solution, (*ii*) autonomously fold chignolin, CLN025, and Trp-cage with folding times that agree with the experimental values, (*iii*) simulate subsequent unfolding and refolding of these miniproteins, and (*iv*) achieve a robust Z score of 1.33 for refining protein models TMR01, TMR04, and TMR07. By comparison, the latest general-purpose AMBER forcefield FF14SB locks the C14-C38 bond to the right-handed configuration in solution under the same protein simulation conditions. Statistical survival analysis shows that FF12MC folds chignolin and CLN025 in isobaric-isothermal MD simulations 2-4 times faster than FF14SB under the same protein simulation conditions. These results suggest that FF12MC may be used for protein simulations to study kinetics and thermodynamics of miniprotein folding as well as protein structure and dynamics.

## INTRODUCTION

Used in computer simulations to describe the relationship between a molecular structure and its energy, an additive (*viz*., nonpolarizable) forcefield is an empirical potential energy function with a set of parameters that is often in the form of Eq. 1.^1–18^ In Eq. 1, *k_b_* and *b_o_* are constants of the bond potential energy for two atoms separated by one covalent bond; *k_θ_* and *θ_o_* are constants of the angle potential energy for two atoms separated by two consecutive covalent bonds; *k_ϕ_* and *δ* are constants of the torsion potential energy for two atoms separated by three consecutive covalent bonds; A_*ij*_ and B_*ij*_ are constants of the van der Waals interaction energy for two intermolecular atoms or for two intramolecular atoms separated by three or more consecutive covalent bonds; C is a constant of the electrostatic interaction energy for two intermolecular atoms or for two intramolecular atoms separated by three or more consecutive covalent bonds. The *A_ij_* and *B_ij_* constants for the atoms separated by three consecutive covalent bonds are typically divided by a 1-4 van der Waals interaction scaling factor (termed SCNB in AMBER forcefields^15,16^). The C constant for the atoms separated by three consecutive covalent bonds is also divided by a 1-4 electrostatic interaction scaling factor (termed SCEE in AMBER forcefields).

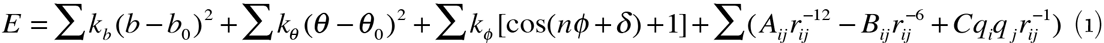

Current general-purpose forcefields are already well refined for various simulations of proteins and other molecules, including folding simulations of a range of miniproteins with implicit or explicit solvation.^12,19–23^ However, simulations using these forcefields to autonomously fold miniproteins in molecular dynamics (MD) simulations with explicit solvation without biasing the simulation systems^21^ have been limited to those performed on extremely powerful but proprietary special-purpose supercomputers.^23–25^ It is desirable to develop a further-refined, special-purpose forcefield that can fold miniproteins with folding times that are both shorter than those using a general-purpose forcefield and, more importantly, closer to the experimental values. This type of specialized forcefield may enable autonomous folding of fast-folding miniproteins in simulations with explicit solvation on commercial computers such as Apple Mac Pros and permit such simulations to be performed under isobaric–isothermal (NPT) conditions that are used in most experimental protein folding studies. It may also enable autonomous folding of slow-folding miniproteins on the special-purpose supercomputers. More importantly, this type of forcefield may improve sampling of nonnative states of a miniprotein in multiple, distinct, independent, unrestricted, unbiased, and classical NPT MD simulations to capture the major folding pathways^26^ and thereby correctly predict the folding kinetics of the miniprotein. It may also improve simulations of genuine localized disorders of folded globular proteins and refinement of comparative models of monomeric globular proteins by MD simulations.^27–44^

It has been shown that uniform reduction of the atomic masses of the entire simulation system (both solute and solvent) by tenfold can enhance configurational sampling in NPT MD simulations.^45^ The uniformly reduced masses by tenfold are hereafter referred to as low masses. The effectiveness of the low-mass NPT MD simulation technique can be explained as follows:^46^ To determine the relative configurational sampling efficiencies of two simulations of the same molecule—one with standard masses and another with low masses, the units of distance [*l*] and energy [*m*]([*l*]/[*t*])^2^ of the low-mass simulation are kept identical to those of the standard-mass simulation, noting that energy and temperature have the same unit. This is so that the structure and energy of the low-mass simulation can be compared to those of the standard-mass simulation. Let superscripts ^lmt^ and ^smt^ denote the times for the low-mass and standard-mass simulations, respectively. Then [*m*^lmt^] = 0.1 [*m*^smt^], [*l*^lmt^] = [*l*^smt^], and [*m*^lmt^]([*l*^lmt^]/[*t*^lmt^])^2^ = [*m*^smt^]([*l*^smt^]/[*t*^smt^])^2^ lead to 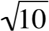 [*t*^lmt^]=[*t*^smt^] A conventional MD simulation program takes the timestep size (Δ*t*) of the standard-mass time rather than that of the low-mass time. Therefore, the low-mass MD simulation at 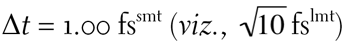 is theoretically equivalent to the standard-mass MD simulation at 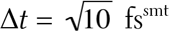, as long as both standard-and low-mass simulations are carried out for the same number of timesteps and there are no precision issues in performing these simulations. This equivalence of mass scaling and timestep-size scaling explains why the low-mass NPT MD simulation at Δ*t* = 1.00 fs^smt^ (*viz*., 3.16 fs^lmt^) offer better configurational sampling efficacy than the standard-mass NPT MD simulation at Δ*t* = 1.00 fs^smt^ or Δ*t* = 2.00 fs^smt^. It also explains why the kinetics of the low-mass simulation can be converted to the kinetics of standard-mass simulation simply through scaling the low-mass time by a factor of 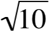. Further, this equivalence explains there are limitations on the use of the mass reduction technique to improve configurational sampling efficiency. Lengthening the timestep size inevitably reduces integration accuracy of an MD simulation. However, the integration accuracy reduction caused by a timestep-size increase is temperature dependent. Therefore, to avoid serious integration errors, low-mass NPT MD simulations must be performed with the double-precision floating-point format and at Δ*t* ≤ 1.00 fs^smt^ and a temperature of ≤340 K.^46^

Because temperatures of biological systems rarely exceed 340 K and because MD simulations are performed typically with the double-precision floating-point format, low-mass NPT MD simulation is a viable configurational sampling enhancement technique for protein simulations at a temperature of ≤340 K.

Another study showed that shortening C-H bonds according to the lengths found in high resolution cryogenic protein structures can reduce the computing time of an MD simulation to capture miniprotein folding.^47^ This is presumably because the shortened C-H bonds reduce the exaggeration of short-range repulsions caused by the implementation of the 6-12 Lennard-Jones potential and a nonpolarizable charge model in an additive forcefield.^48^ A subsequent study found that increasing or decreasing SCNBs of ϕ and ψ and/or SCEEs of ϕ and ψ can raise or lower, respectively, the ratio of the α-helical conformation over the β-strand conformation.^49^ This suggests that the propensities of a forcefield to adopt secondary structure elements can be adjusted by modifying SCNBs and/or SCEEs of ϕ and ψ without implementing the four backbone torsions (ϕ, ψ, ϕ’ and ψ’).

In this context and aiming to simulate proteins in MD simulations with explicit solvation, this author devised an additive forcefield named FF12MCsm that is based on general-purpose AMBER forcefield FF99^50^ with (*i*) the aliphatic C-H bonds shortened to 0.98 Å and the aromatic C-H bonds shortened to 0.93 Å, (*ii*) removal of torsions involving a nonperipheral *sp^3^* atom, and (*iii*) reduced 1-4 interaction scaling factors of torsions *ϕ* and *ψ*(1.00 for SCNB; 1.18 for SCEE). The shortened bond lengths were obtained from a survey of 3709 C-H bonds in the cryogenic protein structures with resolutions of 0.62-0.79 Å.^47^ The reduced scaling factors were obtained from benchmarking FF12MCsm against the experimentally determined mean helix content of Ac-(AAQAA)_3_-NH_2_ (hereafter abbreviated as AAQAA).^51^ To avoid replacing the nonperipheral-*sp*^3^ torsion parameters with a set of arbitrary and complicated scaling factors, two requirements were used to determine the SCNB and SCEE for torsions *ϕ* and *ψ* in FF12MCsm. First, the computed mean α-helix contents of AAQAA at different temperatures using a reported NPT MD simulation protocol^49^ had to be close to the experimental values. Second, the SCNB and SCEE had to be close to 1.00, namely, the scaling of *ϕ* and *ψ* should be reduced as much as possible. As described in RESULTS AND DISCUSSION, with SCNB reduced from 2.00 in FF99 to 1.00 in FF12MCsm and SCEE reduced from 1.20 in FF99 to 1.18 in FF12MCsm, the computed mean α-helix contents of AAQAA using FF12MCsm are indeed close to the experimental data. Like the removal of the 1-4 interaction scaling factors in the GLYCAM06 forcefield,^52^ the scaling of the 1-4 van der Waals interactions for *ϕ* and *ψ* is completely removed in FF12MCsm. The scaling of the 1-4 electrostatic interactions for *ϕ* and *ψ* is also reduced.

Also as demonstrated in RESULTS AND DISCUSSION, these modifications in combination with the mass reduction technique enabled FF12MCsm to fold miniproteins with folding times that were substantially shorter than those of a general-purpose forcefield. However, FF12MCsm did not fold the miniproteins with folding times that were shorter than the experimentally observed folding times, which emphasizes that these modifications were not made to artificially accelerate folding rates for saving computing time. Instead these modifications were made to improve (*i*) sampling of nonnative states of a miniprotein, (*ii*) simulating genuine localized disorders of a folded globular protein, and (*iii*) refining comparative models of a monomeric globular protein.

As reported,^46^ FF12MCsm is intended for standard-mass MD simulations with an explicit solvation model at Δ*t* ≤3.16 fs^smt^ and a temperature of ≤340 K without employing the hydrogen mass repartitioning scheme.^53–55^ FF12MCsm can also be used for standard-mass MD simulations at Δ*t* >3.16 fs^smt^ and a temperature of >340 K with the hydrogen mass repartitioning scheme. A combination of FF12MCsm with the low-mass configurational sampling enhancement technique^45,46^ is a derivative of FF12MCsm. With all atomic masses uniformly reduced by tenfold, this derivative (hereafter abbreviated as FF12MC) is intended for low-mass NPT MD simulations of proteins with an explicit solvation model (preferably the TIP3P water model^56^) at Δ*t* = 1.00 fs^smt^ and a temperature of ≤340 K.

This article reports an FF12MC evaluation study consisting of 1350 NPT MD simulations at 1 atm and 274-340 K with an aggregated simulation time of 1252.572 μs^smt^. Using general-purpose AMBER forcefields FF96 (see **RESULTS AND DISCUSSION** for reasons to include this forcefield),^57^ FF12SB, and FF14SB^16^ as references, these simulations were carried out to determine whether in multiple, distinct, independent, unrestricted, unbiased, and classical NPT MD simulations FF12MC or FFnMCsm can (*i*) reproduce the experimental *J*-coupling constants of four cationic homopeptides (Ala_3_, Ala_5_, Ala_7_, and Val_3_)^58^ and four folded globular proteins of the third immunoglobulin-binding domain of protein G (GB3),^59,60^ bovine pancreatic trypsin inhibitor (BPTI),^61^ ubiquitin,^62^ and lysozme,^63^ (*ii*) reproduce crystallographic B-factors^64^ and nuclear magnetic resonance (NMR)-derived Lipari-Szabo order parameters^65^ of GB3, BPTI, ubiquitin, and lysozyme, (*iii*) simulate the experimentally observed flipping between left-and right-handed configurations for the C14-C38 disulfide bond of BPTI and its mutant,^66^ (*iv*) autonomously fold β-hairpins of chignolin^67^ and CLN025^68^ and an α-miniprotein Trp-cage (the TC10b sequence^69^) with folding times (τ_f_s) in agreement with experimental τ_f_s,^70,71^ (*v*) simulate subsequent unfolding and refolding of these sequences, and (*vi*) refine TMR01, TMR04, and TMR07—comparative models of proteins selected from the first Critical Assessment of protein Structure Prediction model Refinement (CASPR) experiment (http://predictioncenter.org/caspR/, note that subsequent model refinement experiments are called CASP rather than CASPR). Unless otherwise specified, all simulations described below are multiple, distinct, independent, unrestricted, unbiased, and classical NPT MD simulations.

## METHODS

### MD simulations of peptides, miniproteins, and folded globular proteins

A peptide or a miniprotein in a fully extended backbone conformation (or a globular protein in its folded state) was solvated with the TIP3P water^56^ with or without surrounding counter ions and/or NaCls and then energy-minimized for 100 cycles of steepest-descent minimization followed by 900 cycles of conjugate-gradient minimization to remove close van der Waals contacts using SANDER of AMBER 11 (University of California, San Francisco). The resulting system was heated from 0 to a temperature of 274-340 K at a rate of 10 K/ps under constant temperature and constant volume, then equilibrated for 10^6^ timesteps under constant temperature and constant pressure of 1 atm employing isotropic molecule-based scaling, and finally simulated in 20 or 30 distinct, independent, unrestricted, unbiased, and classical NPT MD simulations using PMEMD of AMBER 11 with a periodic boundary condition at 274-340 K and 1 atm. The fully extended backbone conformations (*viz*., anti-parallel β-strand conformations) of Ala_3_, Ala_5_, Ala_7_, Val_3_, AAQAA, chignolin, CLN025, and Trp-cage were generated by MacPyMOL Version 1.5.0 (Schrödinger LLC, Portland, OR). The folded globular protein structures of GB3, BPTI, mutant of BPTI, ubiquitin, and lysozyme were taken from the Protein Data Bank (PDB) structures of IDs 1P7E/1IGD, 5PTI/1PIT, 1QLQ, 1UBQ, and 4LZT, respectively. Four crystallographically determined interior water molecules (WAT111, WAT112, WAT113, and WAT122) were included in the 5PTI structure as the initial conformation of the simulations. Likewise, five interior water molecules (WAT2017, WAT2023, WAT2025, WAT2072, and WAT2092) were included the initial 1QLQ structure. CASPR models TMR01, TMR04, and TMR07 were downloaded from http://predictioncenter.org/caspR/. For TMR01, the *cis* amide bond of Ser70 was manually changed to the *trans* configuration, and all residues that were not determined in the corresponding crystal structure (PDB ID: 1XE1) were removed. His28, His33, His44, and His68 of TMR04 were treated as HIE. His20, His51, and His53 of TMR07 were treated as HID. The numbers of TIP3P waters and surrounding ions, initial solvation box size, ionizable residues, and computers used for the NPT MD simulations are provided in Table S1. The 30 unique seed numbers for initial velocities of Simulations 1-30 are listed in Table S2. All simulations used (*i*) a dielectric constant of 1.0, (*ii*) the Berendsen coupling algorithm,^72^ (*iii*) the Particle Mesh Ewald method to calculate electrostatic interactions of two atoms at a separation of >8 Å,^73^ (*iv*) Δ*t* = 0.10, 1.00, or 3.16 fs^sm^, (*v*) the SHAKE-bond-length constraints applied to all bonds involving hydrogen, (*vi*) a protocol to save the image closest to the middle of the “primary box” to the restart and trajectory files, (*vii*) a formatted restart file, (*viii*) the revised alkali and halide ions parameters,^74^ (*ix*) a cutoff of 8.0 Å for nonbonded interactions, (*x*) the atomic masses of the entire simulation system (both solute and solvent) were either unscaled or reduced uniformly by tenfold, and (*xi*) default values of all other inputs of the PMEMD module. For the simulations of Ala_3_, Ala_5_, Ala_7_, and Val_3_, the forcefield parameters of the cationic Ala (ALC) and the cationic Val (VAC) with their amino and carboxylate groups protonated at pH 2 were generated according to a published procedure using both α-helix and β-strand conformations for the RESP charge calculation.^75^ These forcefield parameters are provided in Supporting Information ALC.lib and VAC.lib. The forcefield parameters of FF12MC are available in the Supporting Information of Ref. ^46^.

### Aggregated native state population

Cα and Cβ root mean square deviation (CαβRMSD) was calculated using PTRAJ of AmberTools 1.5 with root mean square (RMS) fit of all α and β carbon atoms to the corresponding ones in the reference structure without mass weighing. Cα root mean square deviation (CαRMSD) or all-carbon root mean square deviation (CRMSD) was calculated similarly with RMS fit of all α carbon atoms or all carbon atoms to the corresponding ones in the reference structure, respectively.

In NPT MD simulations, chignolin could fold to its native β-hairpin with Tyr2 and Trp9 on the same side of the hairpin^67^ (Fig. 1A) and to native-like β-hairpins with Tyr2 on one side of the hairpin and Trp9 on the other (Fig. 1B).^47^ Similarly, CLN025 could fold to native-like β-hairpins with Tyr1, Trp9, and Tyr10 on one side of the β-sheet and Tyr2 on the other (Fig. 1K) or with Tyr1 and Trp9 on one side and Tyr2 and Tyr10 on the other (Fig. 1L) in NPT MD simulations,^45^ while the native conformations of CLN025 in the NMR and crystal structures have Tyr2 and Trp9 on one side of the β-sheet and Tyr1 and Tyr10 on the other^68^ (Fig. 1F and G).

**Fig. 1.**
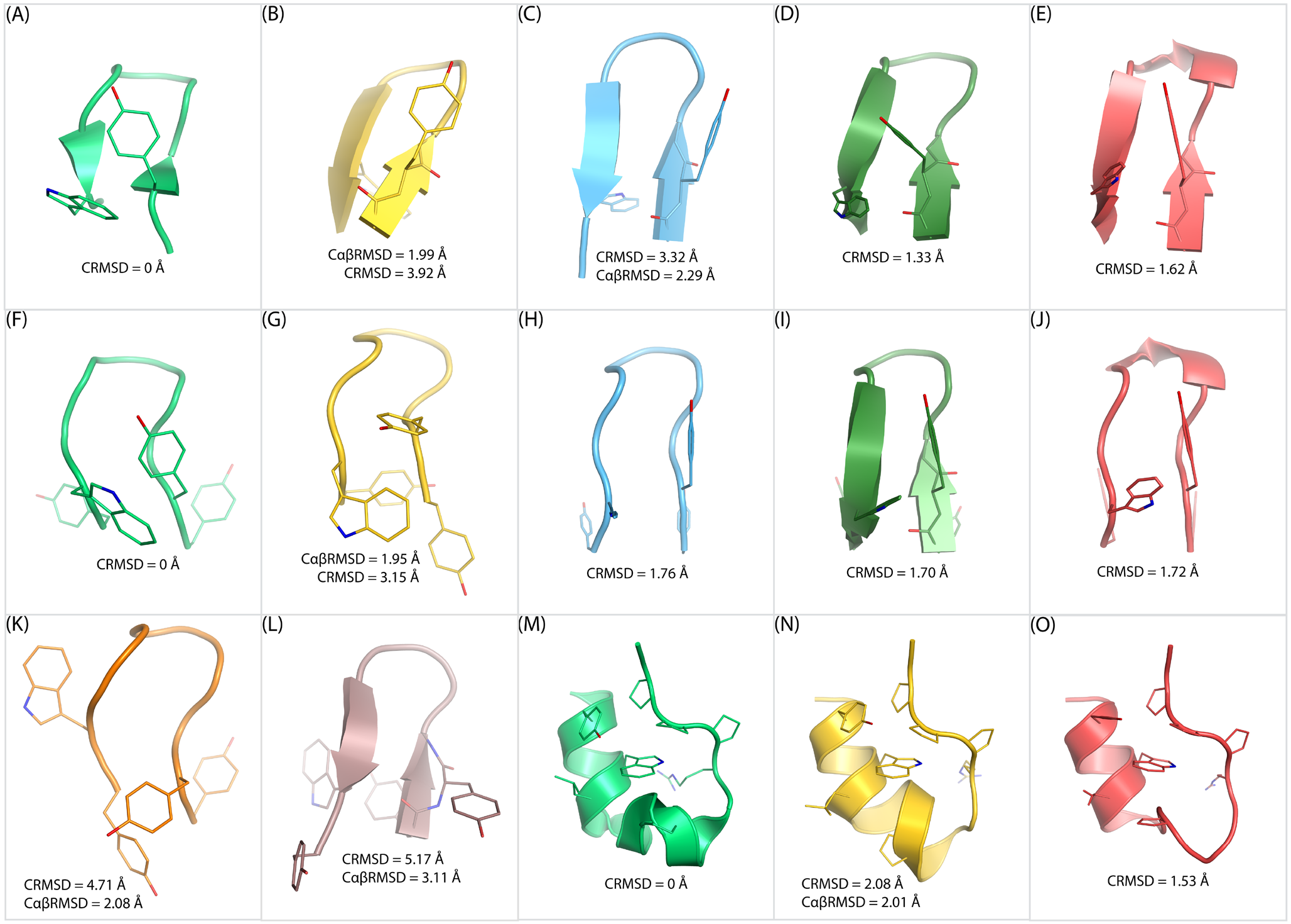
Native and native-like conformations of chignolin, CLN025, and Trp-cage (TC10b). (*A*) The NMR structure of chignolin. (*B*) A native-like conformation of chignolin generated by FF12SB. (*C*) The average chignolin conformation of the largest cluster generated by FF12SB. (*D*) The average chignolin conformation of the largest cluster generated by FF14SB. (*E*) The average chignolin conformation of the largest cluster generated by FF12MC. (*F*) The NMR structure of CLN025. (*G*) The crystal structure of CLN025. (*H*) The average CLN025 conformation of the largest cluster generated by FF12SB. (*I*) The average CLN025 conformation of the largest cluster generated by FF14SB. (*J*) The average CLN025 conformation of the largest cluster generated by FF12MC. (*K*) A native-like conformation of CLN025 generated by FF12SB. (*L*) Another native-like conformation of CLN025 generated by FF12SB. (*M*) The NMR structure of the Trp-cage. (*N*) A native-like conformation of the Trp-cage generated by FF12MC. (*O*) The average Trp-cage conformation of the largest cluster generated by FF12MC.

The smallest CαβRMSD between one of the native-like chignolin conformations and the chignolin NMR structure is 1.99 Å, whereas the corresponding CαRMSD and CRMSD are 1.58Å and 3.92 Å, respectively (Fig. 1B). The smallest CαβRMSD between one of the native-like CLN025 conformations and the CLN025 NMR structure is 2.08 Å, but the corresponding CαRMSD and CRMSD are 1.33 Å and 4.71 Å, respectively (Fig. 1K). The smallest CαβRMSD and CRMSD between the native and native-like conformations of the Trp-cage (TC10b) are 2.01 Å and 2.08 Å, respectively (Fig. 1N). To distinguish conformations at the native state from those at native-like states (Fig. 1B, K, L, and N) or those at nonnative states, in this study the CαβRMSD cutoff was set at 1.96 Å. Although the time series of CαβRMSD from native conformations revealed that AAQAA, chignolin, CLN025, and the Trp-cage can be folded to conformations with CαβRMSDs of ≤1.50 Å (Fig. S1), the CαβRMSD cutoff for the native state was set at 1.96 Å because the CαβRMSD between the NMR and crystal structures of CLN025 is 1.95 Å (Fig. 1G). Otherwise, using a CαβRMSD cutoff of ≤1.50 Å would preclude the conformation determined by the crystallographic analysis that is commonly considered at the native state.

Therefore, the individual native state population of chignolin, CLN025, AAQAA, or the Trp-cage in one MD simulation was calculated as the number of conformations with CαβRMSDs of ≤1.96 Å divided by the number of all conformations saved at every 10^5^ timesteps. Averaging the individual native state populations of a set of 20 or 30 distinct and independent simulations gave rise to the aggregated native state population for the set. The standard deviation (SD) and standard error (SE) of the aggregated native state population were calculated according to Eqs. 1 and 2 of Ref. ^47^, respectively, wherein *N* is the number of all simulations, *P_i_* is the individual native state population of the *i*th simulation, and 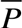 is the aggregated native state population.

### Fractional helicity and α-helix population of AAQAA

**The experimentally determined fractional helicity** (or mean helix content) of AAQAA at a specific temperature (in units of °C) was estimated by averaging component helicities that were obtained according to Eqs. 1 and 2 of Ref. ^51^ with *T*_m_ and Δ*T* values and their SDs taken from Table I of Ref. ^51^. Torsions *ϕ* and *ψ* of each residue in AAQAA were computed from 2 × 10^7^ conformations saved at every 10^3^ timesteps of 20 1.00-μs^smt^ or 3.16-μs^smt^ simulations of AAQAA with the simulation conditions described above. The forcefield parameters for the Ala residue with amidation using NH_2_ (ALN) were taken from Ref. ^49^. **The computationally determined fractional helicity** of AAQAA was calculated from *ϕ* and *ψ* as follows: A residue was considered to be in the α-helical (*viz*., 3.6_13_-helical) conformation if it was one of four consecutive residues with all their torsions *ψ* and *ϕ* within 20° of the reported *ψ* and *ϕ* for α-helix (*ϕ* of −57° and *ψ* of −47°).^76^ A component fractional helicity of a residue in AAQAA was defined as the number of the α-helix conformations for that residue divided by the number of all conformations for AAQAA (*viz*., 2 × 10^7^). Averaging the component fractional helicities of residues 1-15 gave rise to the computationally determined fractional helicity of AAQAA. **The α-helix population** of AAQAA was calculated from CαβRMSD as follows: Cluster analysis of 20,000 conformations from the 20 3.16-μs^smt^ simulations of AAQAA using FF12MC identified a full-α-helix conformation with hydrogen bonds involving the Ac and NH_2_ terminal groups (Fig. 2A) as the most popular conformation (Table S3). Using this conformation as the native conformation, CαβRMSDs for all 2 × 10^7^ conformations of AAQAA were then calculated to determine the number of conformations with CαβRMSDs of ≤1.96 Å. Dividing this number by the number of all AAQAA conformations gave rise to the α-helix population of AAQAA. The SDs of the computationally determined fractional helicity and the α-helix population were calculated using the same method for the SD of the aggregated native state population described above.

**Fig. 2.**
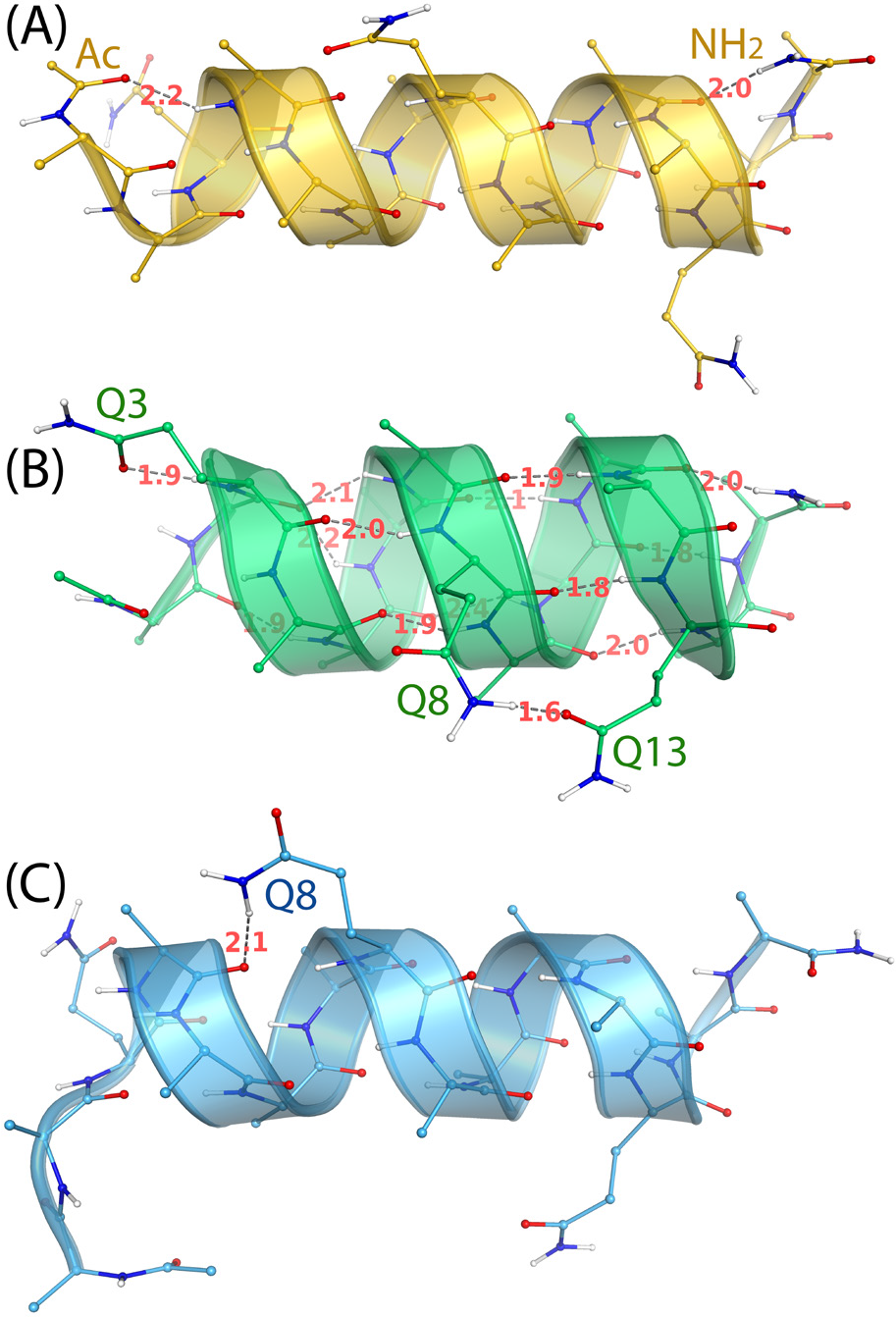
The three most populated, instantaneous conformations of AAQAA observed in MD simulations using FF12MC. Numbers in red denote hydrogen bond lengths in Å. (*A*) The full-α-helix conformation showing hydrogen bonds of two terminal protecting groups. (*B*) The α-and-π helical conformation showing the side-chain • main-chain hydrogen bond of Gln3, the side-chain • side-chain hydrogen bond of Gln8 and Gln13, and main-chain • main-chain hydrogen bonds in α and π helices. (*C*) The α-helix conformation showing substantial unfolding of the Ac, Ala1, and NH_2_ residues.

**Table I.**
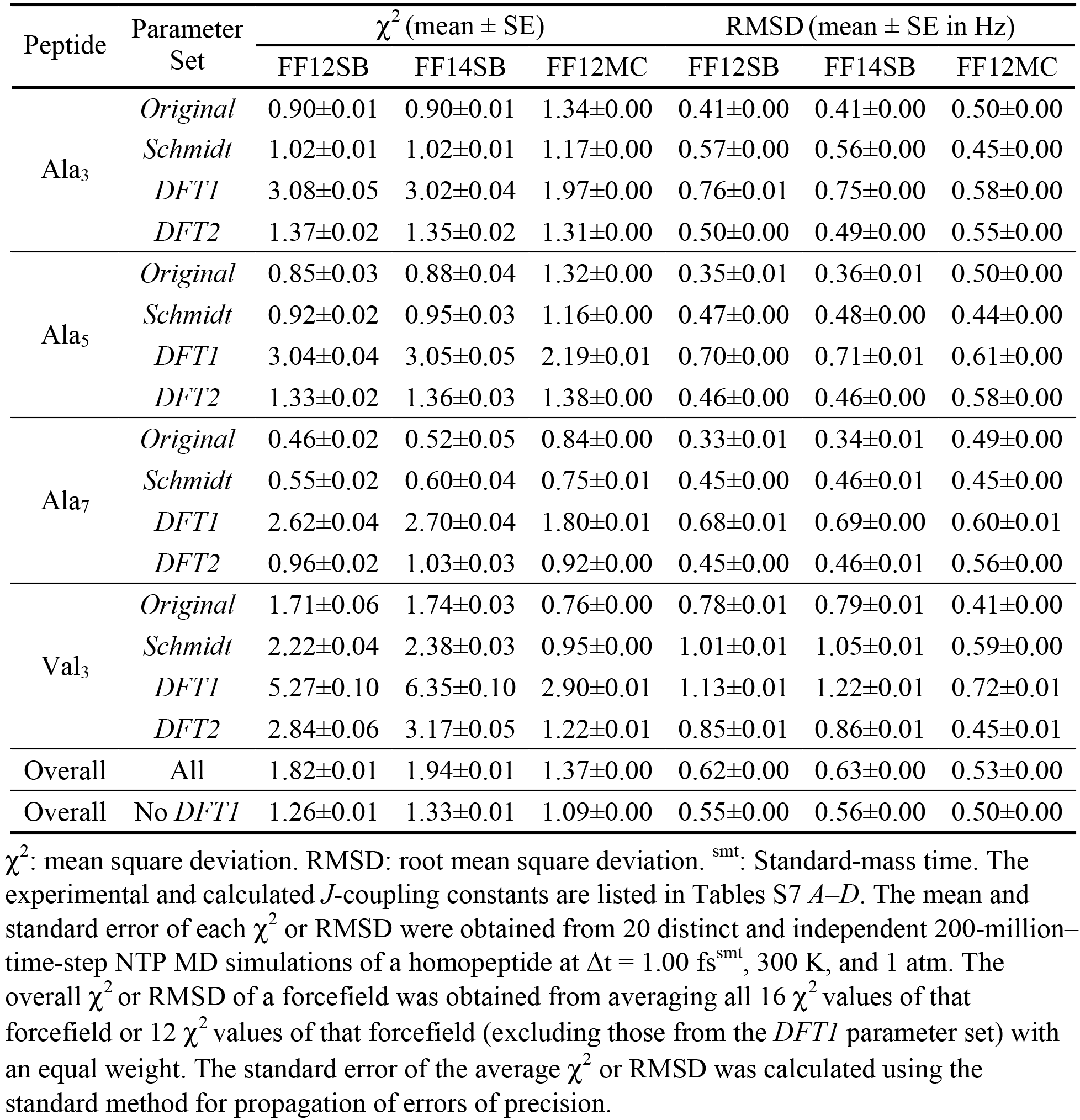
Mean square deviations and root mean square deviation between experimental and calculated *J*-coupling constants of homopeptides using different parameter sets of the Karplus equations.

| Peptide | Parameter Set | $\chi^2$ (mean ± SE) | | | RMSD (mean ± SE in Hz) | | |
| --- | --- | --- | --- | --- | --- | --- | --- |
|  |  | FF12SB | FF14SB | FF12MC | FF12SB | FF14SB | FF12MC |
| Ala<sub>3</sub> | *Original* | 0.90±0.01 | 0.90±0.01 | 1.34±0.00 | 0.41±0.00 | 0.41±0.00 | 0.50±0.00 |
| | *Schmidt* | 1.02±0.01 | 1.02±0.01 | 1.17±0.00 | 0.57±0.00 | 0.56±0.00 | 0.45±0.00 |
| | *DFT1* | 3.08±0.05 | 3.02±0.04 | 1.97±0.00 | 0.76±0.01 | 0.75±0.00 | 0.58±0.00 |
| | *DFT2* | 1.37±0.02 | 1.35±0.02 | 1.31±0.00 | 0.50±0.00 | 0.49±0.00 | 0.55±0.00 |
| Ala<sub>5</sub> | *Original* | 0.85±0.03 | 0.88±0.04 | 1.32±0.00 | 0.35±0.01 | 0.36±0.01 | 0.50±0.00 |
| | *Schmidt* | 0.92±0.02 | 0.95±0.03 | 1.16±0.00 | 0.47±0.00 | 0.48±0.00 | 0.44±0.00 |
| | *DFT1* | 3.04±0.04 | 3.05±0.05 | 2.19±0.01 | 0.70±0.00 | 0.71±0.01 | 0.61±0.00 |
| | *DFT2* | 1.33±0.02 | 1.36±0.03 | 1.38±0.00 | 0.46±0.00 | 0.46±0.00 | 0.58±0.00 |
| Ala<sub>7</sub> | *Original* | 0.46±0.02 | 0.52±0.05 | 0.84±0.00 | 0.33±0.01 | 0.34±0.01 | 0.49±0.00 |
| | *Schmidt* | 0.55±0.02 | 0.60±0.04 | 0.75±0.01 | 0.45±0.00 | 0.46±0.01 | 0.45±0.00 |
| | *DFT1* | 2.62±0.04 | 2.70±0.04 | 1.80±0.01 | 0.68±0.01 | 0.69±0.00 | 0.60±0.01 |
| | *DFT2* | 0.96±0.02 | 1.03±0.03 | 0.92±0.00 | 0.45±0.00 | 0.46±0.01 | 0.56±0.00 |
| Val<sub>3</sub> | *Original* | 1.71±0.06 | 1.74±0.03 | 0.76±0.00 | 0.78±0.01 | 0.79±0.01 | 0.41±0.00 |
| | *Schmidt* | 2.22±0.04 | 2.38±0.03 | 0.95±0.00 | 1.01±0.01 | 1.05±0.01 | 0.59±0.00 |
| | *DFT1* | 5.27±0.10 | 6.35±0.10 | 2.90±0.01 | 1.13±0.01 | 1.22±0.01 | 0.72±0.01 |
| | *DFT2* | 2.84±0.06 | 3.17±0.05 | 1.22±0.01 | 0.85±0.01 | 0.86±0.01 | 0.45±0.01 |
| Overall | All | 1.82±0.01 | 1.94±0.01 | 1.37±0.00 | 0.62±0.00 | 0.63±0.00 | 0.53±0.00 |
| Overall | No *DFT1* | 1.26±0.01 | 1.33±0.01 | 1.09±0.00 | 0.55±0.00 | 0.56±0.00 | 0.50±0.00 |

### *J*-coupling constant calculation

Using PTRAJ of AmberTools 1.5, torsions *ϕ* and *ψ* of each residue in a homopeptide were computed from all conformations saved at every 10^3^ timesteps of 20 simulations of the peptide with the simulation conditions described above. Similarily, torsions *ϕ* and *ψ* of each residue and torsion χ of each non-glycine residue in a folded globular protein were computed from all conformations saved at every 10^5^ timesteps of 20 simulations of the protein. An instant *J*-coupling constant (*J_i_* in Hz) of a residue was calculated according to Eqs. S1-S20 using a set of parameters described as follows. The *Original* parameters of Eqs. S1-S5, S6, S7, and S8 were taken from Refs. ^62, 77, 78^, and ^79^, respectively. The *Schmidt* parameters of Eqs. S1-S5, S6, S7, and S8 were taken from Refs. ^80^, ^77^, ^78^, and ^79^, respectively. The DFT1 and DFT2 parameters of Eqs. S1-S5, S6, S7, and S8 were taken from Refs. ^81^, ^77^, ^78^, and ^79^, respectively. The *Original* and *Schmidt* parameters of Eqs. S9-S14 were taken from Ref. ^82^. The *Best-Fit* and DFT parameters of Eqs. S15-S20 were taken from Ref. ^83^. Averaging all instant *J*-coupling constants of a residue gave rise to the *J*-coupling constant for that residue.

The mean square deviation (χ^2^) between experimental and calculated *J*-coupling constants was estimated according to Eq. S21 with σ_i_ values taken from Table S3 of Ref. ^84^. The mean and SE of a χ^2^ value were obtained from 20 simulations using the same method for the mean and SE of the aggregated native state population described above. The experimental *J*-coupling constants of the four homopeptides were obtained from the supporting information of Ref. ^58^. The experimental *J*-coupling constants of the four folded globular proteins were obtained from the supporting information of Ref. ^17^ for GB3 and ubiquitin, Ref. ^85^ for BPTI, and Refs. ^63^ and ^85^ for lysozyme. The simulation temperatures of the protein *J*-coupling constant calculations were taken from Refs. ^59^ and ^60^ for GB3, Ref. ^61^ for BPTI, Ref. ^62^ for ubiquitin, and Ref. ^63^ for lysozyme.

The overall χ^2^ value of a forcefield for peptide *J*-coupling constants was obtained by averaging all 16 χ^2^ values of that forcefield in Table I or 12 χ^2^ values of that forcefield in Table I (excluding those using the DFT1 parameter set) with an equal weight. Similarily, the overall χ^2^ value of a forcefield for protein *J*-coupling constants was obtained from averaging all four combined or main-chain χ^2^ values of the forcefield with an equal weight. The SE of the overall χ^2^ was calculated according to the standard method for propagation of errors of precision.^86^

### The Lipari-Szabo order parameter prediction

Using a two-step procedure and PTRAJ of AmberTools 1.5, the backbone N-H Lipari-Szabo order parameter (S^2^)^65^ of a folded globular protein was predicted from all conformations saved at every 10^3^ timesteps of 20 simulations of the protein with the simulation conditions described above and additional conditions described in RESULTS AND DISCUSSION. The first step was to align all saved conformations onto the first saved one using RMS fit of all CA, C, N, and O atoms. The second step was to compute S^2^ using the isotropic reorientational eigenmode dynamics (iRED) analysis method^87^ implemented in PTRAJ. Although the first step was unnecessary for the iRED analysis method,^87^ the explicit alignment was done in this study for the future use of these conformations to compute S^2^ with other analytical methods. PDB IDs 1P7E, 5PTI, 1UBQ, and 4LZT were used in the GB3, BPTI, ubiquitin, and lysozyme simulations to calculate their S^2^ parameters. The temperatures of the simulations for GB3, BPTI, ubiquitin, and lysozyme were set at 297 K, 298 K, 300 K, and 308 K, respectively, according to the temperatures at which the experimental S^2^ parameters were obtained.^88–91^ The calculated S^2^ parameters of each protein reported in Table S4 and Fig. 3 are the average of all S^2^ parameters derived from 20 distinct and independent simulations of the protein. The SE of an S^2^ parameter was calculated using the same method as the one for the SE of an aggregated native state population. The ability of a forcefield to reproduce the experimental S^2^ parameters is determined by root mean square deviation (RMSD) between computed and experimental S^2^ parameters. The experimental S^2^ parameters extracted from ^15^N spin relaxation data for GB3, ubiquitin, lysozyme, and BPTI were obtained from respective supporting information or corresponding authors of Refs. ^88–91^. The SE of an RMSD was calculated using the same method as the one for the SE of an S^2^ value.

**Fig. 3.**
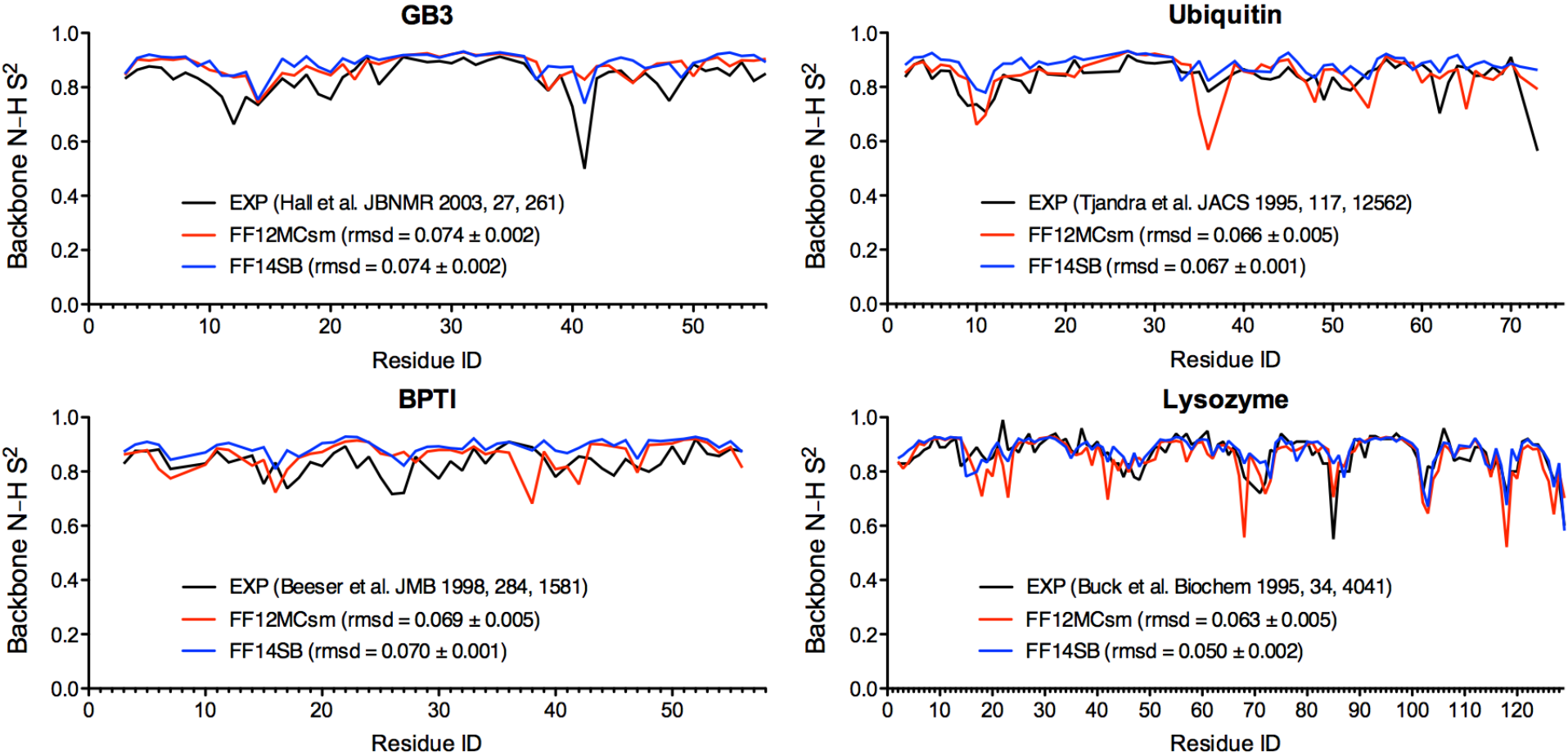
Experimental and calculated Lipari-Szabo order parameters of backbone N-H bonds of GB3, BPTI, ubiquitin, and lysozyme. The order parameters were calculated from 20 unbiased, unrestricted, distinct, independent, and 50-ps^smt^ NPT MD simulations using FF12MCsm or FF14SB with Δ*t* = 0.1 fs^smt^.

### The crystallographic B-factor prediction

Using a two-step procedure and PTRAJ of AmberTools 1.5, the crystallographic B-factors of Cα and Cγ in a folded globular protein were estimated from all conformations saved at every 10^3^ timesteps of 20 simulations of the protein with the simulation conditions described in the Lipari-Szabo order parameter prediction. The first step was to align all saved conformations onto the first saved one to obtain an average conformation using RMS fit of all CA atoms (for Cα B-factors) or all CG and CG2 atoms (for Cγ B-factors). The second step was to RMS fit all CA atoms (or all CG and CG2 atoms) of all saved conformations onto the corresponding atoms of the average conformation and then calculate the Cα (or Cγ) B-factors using the “atomicfluct” command in PTRAJ. PDB IDs 1IGD, 1PIT, 1UBQ, and 4LZT were used in the GB3, BPTI, ubiquitin, and lysozyme simulations to calculate their B-factors. A truncated 1IGD structure (residues 6-61) was used for the GB3 simulations. The simulations for GB3, BPTI, and ubiquitin were done at 297 K, whereas the simulations of lysozyme were performed at 295 K. The calculated B-factors of each protein reported in Table S5 and Fig. 4 are the average of all B-factors derived from 20 distinct and independent simulations of the protein. The SE of a B-factor was calculated using the same method as the one for the SE of an S^2^ parameter. The ability of a forcefield to reproduce the B-factors was measured by RMSD between computed and experimental B-factors. The experimental B-factors of GB3, BPTI, ubiquitin, and lysozyme were taken from the crystal structures of PDB IDs of 1IGD, 4PTI, 1UBQ, and 4LZT, respectively. The SE of an RMSD was calculated using the same method for the SE of a B-factor.

**Fig. 4.**
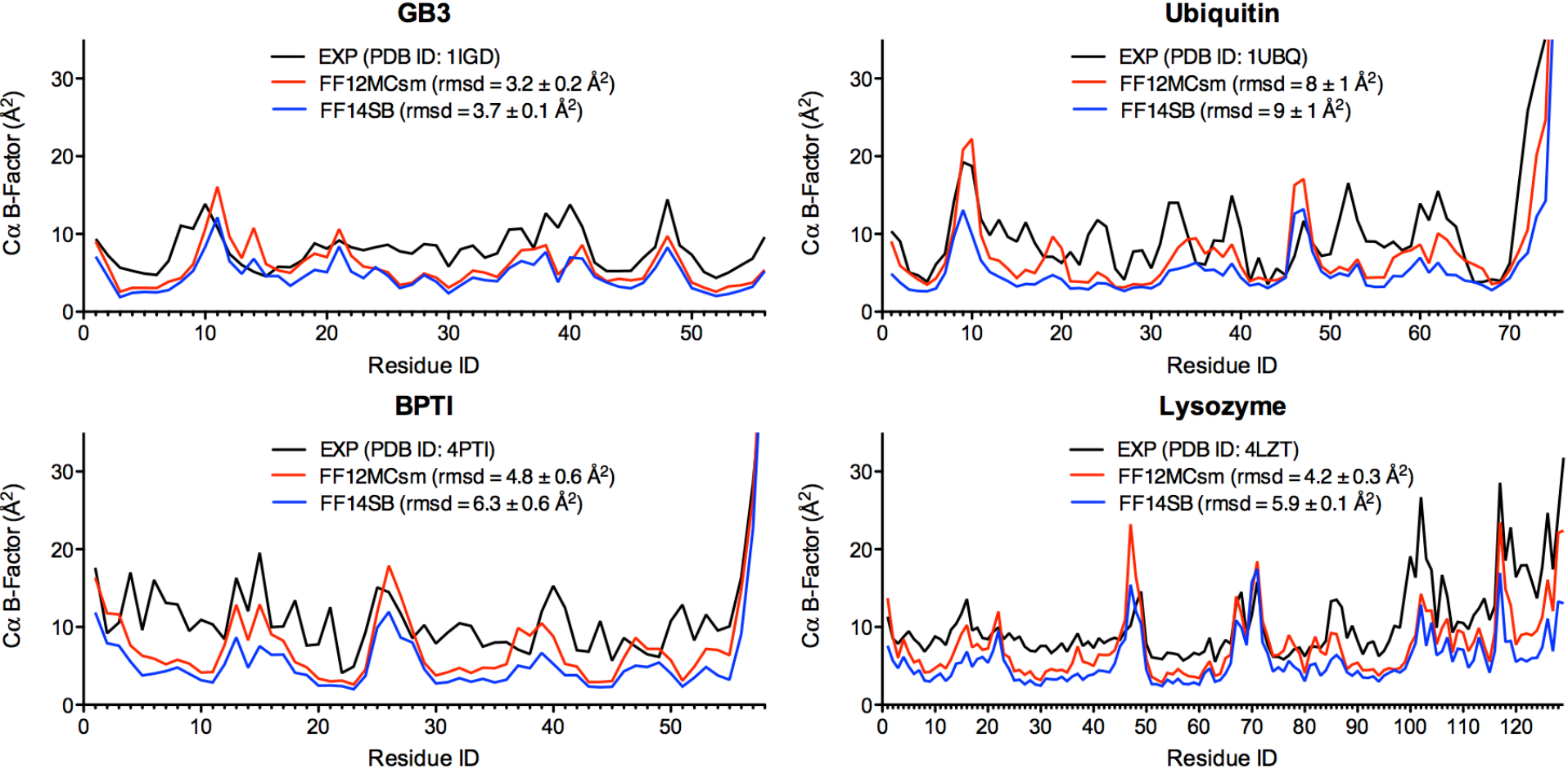
Experimental and calculated crystallographic Cα B-factors of GB3, BPTI, ubiquitin, and lysozyme. The B-factors were calculated from 20 unbiased, unrestricted, distinct, independent, and 50-ps^smt^ NPT MD simulations using FF12MCsm or FF14SB Δ*t* = 0.1 fs^smt^.

### Folding time estimation

The folding time (τ_f_) of a peptide or miniprotein was estimated from the mean time of the peptide or miniprotein to fold from a fully extended backbone conformation to its native conformation (abbreviated hereafter as mean time-to-folding) in 20 (for AAQAA and β-hairpins) or 30 (for the Trp-cage) distinct and independent NPT MD simulations using survival analysis methods^92^ implemented in the R survival package Version 2.38-3 (http://cran.r-project.org/package=survival). The afore-described CαβRMSD cutoff of ≤1.96 Å was used to identify the native conformation. For each simulation with conformations saved at every 10^5^ timesteps, the first time instant at which CαβRMSD reached ≤1.96 Å was recorded as an individual folding time (IFT; Fig. S1). Using the Kaplan-Meier estimator^93,94^ [the Surv() function in the R survival package], the mean time-to-folding was calculated from a set of simulations each of which captured a folding event. If a parametric survival function mostly fell within the 95% confidence interval (95% CI) of the Kaplan-Meier estimation for a set of simulations each of which captured a folding event, the parametric survival function [the Surreg() function in the R survival package] was then used to calculate the mean time-to-folding of that set of simulations. If the mean time-to-folding derived from the Kaplan-Meier estimator for a first set of simulations—each of which captured a folding event—was nearly identical to the one derived from a parametric survival function for the first set, the parametric function was then used to calculate the mean time-to-folding of a second set of simulations that were identical to the first set except that the simulation time or forcefield of the second set was changed. When a parametric survival function was used to calculate the mean time-to-folding, not all simulations in a set had to capture a folding event, but more than half of the set must capture a folding event to avoid an overly wide 95% CI.

### CASPR model refinement evaluation and forcefield performance ranking

The average conformation of the largest cluster of a protein model—identified by the cluster analysis described below—was used as the refined model of the protein. This refined model was evaluated with nine quality scores (QSs) including the sseRMSD score,^37^ the CαRMSD score, the GDT-TS and GDT-HA scores,^95^ the GDC-all score,^96^ the RPF score,^97^ the LDDT score,^98^ the SphereGrinder score,^99^ and the CAD score.^100^ The sseRMSD score was calculated using PTRAJ of AmberTools 1.5 with RMS fit of the CA, C, N, and O atoms of selected residues in the refined model to the corresponding ones in the crystal structure without mass weighing, wherein the selected residues in the refined model correspond to those defined as secondary structure elements in the crystal structure. The CαRMSD, GDT-TS, and GDT-HA scores were calculated using the TM-score program.^101^ The GDC-all score was calculated using the input of “LGA_49605-gdc” at the LGA^102^ server (http://proteinmodel.org/AS2TS/LGA/lga.html). The RPF score was calculated using the RPF program (for Mac OS X) modified for the assessment of template-based protein structure predictions in the 10^th^ Critical Assessment of protein Structure Prediction (CASP10).^97^ This modified program was obtained from Dr. Yuanpen J. Huang of the Gaetano T. Montelione group. The LDDT score was calculated using the LDDT executable (for Mac OS X) downloaded from http://swissmodel.expasy.org/lddt/downloads/. The SphereGrinder score was calculated using the SphereGrinder server (http://spheregrinder.cs.put.poznan.pl). The CAD score was calculated with the all-atom option for both target and model structures using the CAD score server (http://bioinformatics.ibt.lt/cad-score/).

### Cluster analysis and data processing

The conformational cluster analysis of a peptide or miniprotein was performed using PTRAJ of AmberTools 1.5 with the average-linkage algorithm,^103^ epsilon of 2.0 Å, and root mean square coordinate deviation on all Cα and Cβ atoms for AAQAA, chignolin, CLN025, and the Trp-cage. Similarily, the analysis of a folded globular protein was done with root mean square coordinate deviation on Cα atoms of all residues of GB3, BPTI, ubiquitin, and lysozyme or on Cα atoms of residues 9-91 for TMR01, residues 7-70 for TMR04, and residues 1-107 for TMR07 (for additional information see Tables S3 and S6). The torsional cluster analyses for BPTI and its mutant were conducted as follows. Using the PTRAJ program, a set of five consecutive torsions of the C14-C38 bond was calculated from each conformation saved at every 10^5^ timesteps from 20 distinct and independent simulations. The five torsions were defined as (*i*) :14@N :14@CA :14@CB :14@SG; (*ii*) :14@CA :14@CB :14@SG :38@SG; (*iii*) :14@CB :14@SG :38@SG :38@CB; (*iv*) :14@SG :38@SG :38@CB :38@CA; (*v*) :38@SG :38@CB :38@CA :38@N. Each set of these torsions was then compared to all other sets using the criterion that two torsion sets are different if one of the five torsions in one set differs by 60 degrees of arc or more from the corresponding one in the other set. The number of torsion sets in a cluster divided by all torsion sets gave rise to the occurrence of the cluster. No energy minimization was performed on the average conformation of any cluster. Radius of gyration was calculated using PTRAJ of AmberTools 1.5. Smoothed time series of CαβRMSD were generated by PRISM of GraphPad Software (La Jolla, California) using 32 neighbors on each size and 6^th^ order of the smoothing polynomial.

## RESULTS AND DISCUSSION

### Use of different timestep sizes for forcefield evaluation

It was reported that unless the atomic masses of the entire simulation system (both solute and solvent) were reduced uniformly by tenfold, FF14SB was unable to fold CLN025 in 10 500-ns^smt^ simulations at 277 K and Δ*t* = 1.00 fs^smt^.^45^ The ability of FF14SB to fold CLN025 in the low-mass simulations is attributed to the use of a long timestep size (Δ*t* = 3.16 fs^lmt^) in the low-mass simulations, which is due to the equivalence of mass scaling and timestep-size scaling as explained in INTRODUCTION. Because of this equivalence, the integration accuracy of a low-mass simulation at Δ*t* = 1.00 fs^smt^ (*viz*., a standard-mass simulation at Δ*t* = 3.16 fs^smt^) can be assumed to be lower than that of a standard-mass simulation at Δ*t* = 1.00 fs^smt^. According to a theoretical analysis^53^ and a study with 160 submicrosecond or microsecond simulations to autonmously fold β-hairpins at different Δ*t*s and different temperatures,^46^ Δ*t* = 3.16 fs^lmt^ for low-mass simulations (or 3.16 fs^smt^ for standard-mass simulations) is still below the integration step size that can cause fatal integration errors as long as the simulations are performed at a temperature of ≤340 K. Informed with this background information, to compare FF12MC with FF12SB/FF14SB, standard-mass simulations with FF12SB/FF14SB and Δ*t* = 1.00 fs^smt^ were used for peptides and miniproteins. This was so that the integration accuracy of such simulations is higher than that of the low-mass simulations with FF12MC and Δ*t* = 3.16 fs^smt^. Low-mass simulations with FF14SB and Δ*t* = 3.16 fs^lmt^ were used only for proteins or in limited cases for miniproteins for direct comparison to low-mass simulations with FF12MC and Δ*t* = 3.16 fs^lmt^.

### Effect of Δ*t* = 3.16 fs^smt^ on quality of NPT MD simulations at a temperature of ≤340 K

As a measure of the integration accuracy or the quality of an MD simulation, 〈Δ*E*^2^〉^1/2^/〈Δ*KE*^2^〉^1/2^ is the ratio of the square root of the fluctuations in the total energy of the simulation system to the square root of the fluctuations in the kinetic energy of the system; the lower the ratio the higher the simulation quality.^104,105^ Although Δ*t* = 3.16 fs^smt^ for the standard-mass simulations (or Δ*t* = 3.16 fs^lmt^ for the low-mass simulations) is below the limit to cause serious integration errors for an NPT MD simulation that uses a thermostat to keep the temperature of the simulation system at a desired value (≤340 K) and remove the accumulated energy caused by integration errors from the system to the thermostat,^46^ Δ*t* = 3.16 fs^smt^ (or Δ*t* = 3.16 fs^lmt^) may still be too long and hence compromise the quality of the NPT simulation. To address this concern, the 〈Δ*E*^2^〉^1/2^/〈Δ*KE*^2^〉^1/2^ ratios were calculated from all NPT simulations described below to compare the integration accuracy of low-mass NPT simulations using FF12MC and Δ*t* = 3.16 fs^lmt^ to that of standard-mass NPT simulations using FF12SB/FF14SB and Δ*t* = 1.00 fs^smt^, noting that the 〈Δ*E*^2^〉^1/2^/〈Δ*KE*^2^〉^1/2^ ratios of the low-mass microcanonical (NVE) MD simulations with FF12MC and Δ*t* = 3.16 fs^lmt^ were not calculated because FF12MC is intended for low-mass NPT MD simulations. It has been reported that all MD simulations carried out to validate FF14SB used Δ*t* = 1.00 or 2.00 fs^smt^, a cutoff of 8.0 Å for nonbonded interactions, and the Particle Mesh Ewald method to calculate electrostatic interactions of two atoms at separations of >8.0 Å.^1^ If the same protocol were used to calculate nonbonded interactions and if the 〈Δ*E*^2^〉^1/2^/〈Δ*KE*^2^〉^1/2^ ratios of the low-mass simulations using FF12MC and Δ*t* = 316 fs^lmt^ were comparable to those of the standard-mass simulations using FF12SB/FF14SB and Δ*t* = 1.00 fs^smt^, it would be reasonable to suggest that Δ*t* = 3.16 fs^smt^ (or Δ*t* = 3.16 fs^lmt^) would not compromise the quality of the NPT MD simulations. Indeed, Table S7 shows that the 〈Δ*E*^2^〉^1/2^/〈Δ*KE*^2^〉^1/2^ ratios (in mean ± SE) of all low-mass NPT simulations using FF12MC at Δ*t* = 3.16 fs^lmt^ range from 0.2405±0.0004 to 0.3685±0.0032, whereas the corresponding ratios of all standard-mass NPT simulations using FF14SB at Δ*t* = 1.00 fs^smt^ range from 0.4097±0.0012 to 0.5064±0.0009. Further, the ranges of the ratio change to 0.3036±0.0008-0.3501±0.0042 and 0.4945±0.0014-0.5011±0.0013 for low-mass NPT simulations using FF14SBlm at At = 3.16 fs^lmt^ and standard-mass NPT simulations using FFnMCsm at Δ*t* = 1.00 fs^smt^, respectively. These data suggest that the use of Δ*t* = 3.16 fs^smt^ at a temperature of ≤340 K does not compromise the NPT MD simulation quality. However, it is not recommended to use Δ*t* > 3.16 fs^smt^ (such as Δ*t* = 4.00 fs^smt^) at a temperature of ≤340 K without employing the hydrogen mass repartitioning scheme^53–55^ because the quality of an MD simulation under such conditions has not been evaluated.

### Reproducing experimental *J*-coupling constants

***J*-coupling constants of homopeptides.** Although it is debatable whether an agreement between experimental and calculated *J*-coupling constants may be used as an indicator of the goodness of fit of a forcefield,^106^ testing the ability of a forcefield to reproduce experimental *J*-coupling constants has become part of a forcefield evaluation study.^13–17^ While the experimental *J*-coupling constants of cationic homopeptide Ala_5_ were used in parameterizing FF12SB and FF14SB,^16^ no experimental *J*-coupling constants of any cationic homopeptides or folded globular proteins were used to develop FF12MC. How well FF12MC can reproduce the experimental *J*-coupling constants relative to those of FF12SB and FF14SB is important to the critical evaluation of FF12MC. This is because the removal of torsions involving a nonperipheral *sp^3^* atom in FF12MC—a radical difference between FF12MC and general-purpose AMBER forcefields— may impair the ability of FF12MC to reproduce the experimental *J*-coupling constants. Accordingly, a *J*-coupling constant calculation study was carried out to investigate the ability of FF12MC to reproduce the experimental *J*-coupling constants of four cationic homopeptides (Ala_3_, Ala_5_, Ala_7_, and Val_3_) at pH 2^58^ relative to those of FF12SB and FF14SB. Homopeptide Gly_3_ was excluded in this study because a limited data set was used in some of the Karplus parameterizations.^58^

In general, results derived from fewer than 20 simulations are considered unreliable.^107,108^ Therefore, in this study 20 distinct and independent simulations at 300 K and 1 atm were carried out for each of the four homopeptides. The calculated main-chain *J*-coupling constants of each peptide—^3^J(H_N_,Hα), ^3^J(H_N_,C’), ^3^J(Hα,C’), ^3^J(C’,C’), ^3^J(H_N_,Cβ), ^1^J(N,Cα), ^2^J(N,Cα), ^3^J(H_N_,Cα)—are listed in Table S8. Plotting the mean square deviation (χ^2^) between experimental and calculated *J*-coupling constants over logarithm of number of timesteps suggests that χ^2^ values of all four peptides are converged after ten million timesteps for FF12MC, FF12SB, FF14SB (Fig. S2).

When the main-chain *J*-coupling constants were calculated using the original parameters of the Karplus equations (the *Original* parameter set in Eqs. S1-S8^62,77–79^), FF12SB and FF14SB reproduced the alanine constants better than FF12MC, whereas FF12MC reproduced the valine constants better than FF12SB and FF14SB (Table I). Overall, the χ^2^ values ± SEs of FF12MC, FF12SB, and FF14SB are ≤1.34±0.00, ≤1.71±0.06, and ≤1.74±0.03, respectively. The χ^2^ values of FF12SB and FF14SB increased uniformly when alternative parameters of the Karplus equations (the *Schmidt, DFT1, and DFT2* parameter sets in Table S9) were used to calculate the *J*-coupling constants. For FF12MC, the χ^2^ values increased uniformly only when the *DFT1* parameter set was used in the calculation.

Doubling the simulation time for each of the 20 Val_3_ simulations using FF14SB did not reduce the χ^2^ values (Table S10). Repeating the Val_3_ simulations using FF14SB and FF12MC with a cutoff of 9.0 Å for nonbonded interactions and the Particle Mesh Ewald method to calculate electrostatic interactions between atoms at separations of >9.0 Å resulted in χ^2^s that were statistically identical to those with the cutoff of 8.0 Å (Fig. S2 and Table S10). The χ^2^ values and their SEs of FF14SB for Ala_5_ in this study (0.88±0.04 for *Original*; 3.05±0.05 for DFT1; 1.36±0.03 for DFT2; Table I) are consistent with the corresponding χ^2^ values (0.89±0.04 for *Original*; 2.71±0.15 for *DFT1*; 1.22±0.03 for *DFT2*) reported in Tables 1-3 of Ref. ^16^. The overall χ^2^ values of FF12MC, FF12SB, and FF14SB are 1.37±0.00, 1.82±0.01, and 1.94±0.01, respectively. These overall χ^2^ values are reduced to 1.09±0.00, 1.26±0.01, and 1.33±0.01, respectively, when the *DFT1* dataset is excluded. These results show that FF12MC is on par with FF12SB and FF14SB in reproducing main-chain *J*-coupling constants of the four peptides, despite the removal of torsions involving a nonperipheral *sp*^3^ atom in FF12MC.

***J*-coupling constants of folded globular proteins.** Before extending the *J*-coupling constant calculation from short peptides to folded globular proteins, it is worth noting that the proton resonance broadening effect of the proteins is substantially greater than that of the peptides and all cross-peaks involving this resonance are overlapped with other peaks. So ambiguity in assigning protein *J*-coupling constants is inevitable. For example, there are two sets of *J*-coupling constants of GB3, a 56-residue protein with near-perfect assignments of *J*-coupling constants.^17,109^ In Ref. ^17^, ^3^J(Hα,Hβ2) and ^3^J(Hα,Hβ3) are 3.99 and 2.13 for Asp22 and 7.15 and 7.92 for Gln35, respectively. In Ref. ^109^, ^17^, ^3^J(Hα,Hβ2) and ^3^J(Hα,Hβ3) are 2.13 and 3.99 for Asp22 and 7.92 and 7.15 for Gln35, respectively. The discrepancies between these datasets that were independently compiled by two well-respected groups underscore the challenge of assigning *J*-coupling constants without ambiguity. It is also worthy of noting that experimental *J*-coupling constants are averaged on a millisecond timescale,^110^ but MD simulations of folded globular proteins are currently limited to the sub-microsecond or microsecond timescale. Despite these challenges, testing the ability of a forcefield to reproduce protein *J*-coupling constants has also become part of a forcefield evaluation study.^13–17^ To compare FF12MC to FF14SB, the main-chain and side-chain *J*-coupling constants of GB3, BPTI, ubiquitin, or lysozyme were calculated as functions of torsions *ϕ,ψ*, and *χ* that were determined from 20 316-ns^smt^ simulations using the two forcefields. All calculated *J*-coupling constants of the four proteins are listed in Table S11.

When the *Original* parameter sets (Table S9) were used to calculate the main-chain and side-chain *J*-coupling constants, FF12MC and FF14SB reproduced the protein *J*-coupling constants with overall χ^2^s (mean ± SE) of ≤78.0±0.1 and ≤71.5±0.1 for main-chain and/or side-chain constants of the four proteins and overall χ^2^s (mean ± SE) of ≤3.57±0.04 and ≤1.16±0.01 for the main-chain constants, respectively (Table II). FF14SB performs markedly better than FF12MC in reproducing the main-chain *J*-coupling constants of folded globular proteins. According to the overall RMSDs between experimental and calculated constants of the four proteins, FF14SB also performs significantly better than FF12MC in reproducing the main-chain *J*-coupling constants (Table II). The same conclusion could be reached when other parameter sets (Table S9) were used, although the overall χ^2^s and RMSDs of the other parameter sets were larger than those of the *Original* parameter sets. Adding harmonic motion to the Karplus relation for spin-spin coupling^111^ led to the same conclusion, although it slightly improved the χ^2^s and RMSDs. These results demonstrate that FF14SB outperforms FF12MC in reproducing the *J*-coupling constants of the four proteins (Table II). Given the challenges in reproducing protein *J*-coupling constants described above, the relatively poor performance of FF12MC is not sufficient to invalidate FF12MC. Nevertheless, it calls for a further evaluation of the ability of FF12MC to simulate structure and dynamics of the four folded globular proteins.

**Table II.** Mean square deviations and root mean square deviation between experimental and calculated *J*-coupling constants of folded globular proteins using the original parameters of the Karplus equations.

| Protein<br>Temp | Type of $J$ | $\chi^2 \pm \text{SE}$ | | RMSD $\pm \text{SE}$ (Hz) | |
| --- | --- | --- | --- | --- | --- |
|  |  | FF12MC | FF14SB | FF12MC | FF14SB |
| GB3<br>298 K | Main-chain | 2.01±0.02 | 1.09±0.02 | 0.94±0.00 | 0.66±0.01 |
| | Side-chain | 59.0±0.3 | 56.7±0.1 | 2.32±0.00 | 2.25±0.00 |
| | Combined | 18.78±0.08 | 17.43±0.04 | 1.49±0.00 | 1.34±0.00 |
| BPTI<br>309 K | Main-chain | — | — | — | — |
| | Side-chain | 167.63±0.07 | 159.4±0.4 | 4.01±0.00 | 3.91±0.00 |
| | Combined | 167.63±0.07 | 159.4±0.4 | 4.01±0.00 | 3.91±0.00 |
| Ubiquitin<br>303 K | Main-chain | 4.0±0.1 | 1.04±0.02 | 1.18±0.01 | 0.67±0.01 |
| | Side-chain | 48.8±0.2 | 36.9±0.2 | 2.15±0.00 | 1.84±0.00 |
| | Combined | 21.3±0.1 | 14.95±0.07 | 1.63±0.01 | 1.26±0.00 |
| Lysozyme<br>308 K | Main-chain | 4.7±0.1 | 1.34±0.01 | 1.98±0.02 | 1.05±0.00 |
| | Side-chain | 149.4±0.5 | 135.8±0.2 | 4.10±0.01 | 3.94±0.00 |
| | Combined | 104.4±0.4 | 94.0±0.2 | 3.58±0.01 | 3.32±0.00 |
| Overall | Combined | 78.0±0.1 | 71.5±0.1 | 2.68±0.00 | 2.46±0.00 |
| Overall | Main-chain | 3.57±0.05 | 1.16±0.01 | 1.37±0.01 | 0.79±0.00 |

### Simulating folded globular protein structures

**Radius of gyration and CαRMSD from crystal structure.** Given the weak performance of FF12MC in reproducing main-chain protein *J*-coupling constants, it is reasonable to suspect that FF12MC may not be able to simulate structure and dynamics of folded globular proteins. To address this concern, 20 316-ns^smt^ simulations of GB3 were carried out using FF12MC or FF14SB. These simulations used the crystal structure of PDB ID 1IGD as the initial conformation and were performed at 297 K at which the NMR study was conducted for determining the Lipari-Szabo order parameters.^88^ The average of 20,000 conformers of GB3 saved at 100-ps^smt^ intervals of the 20 simulations using FF12MC has a CαRMSD of 0.84 Å relative to the crystal structure, while the corresponding CαRMSD of FF14SB is 0.89 Å (Table III). The mean, SD, and SE of the radius of gyration of the 20,000 GB3 conformers obtained from the 20 simulations using FF12MC are 10.85 Å, 0.11 Å, and 0.02 Å, respectively; while the corresponding ones of FF14SB are 10.97 Å, 0.11 Å, and 0.02 Å, respectively (Table III). By comparison, the CαRMSD of the average of 60 NMR conformers is 0.80 Å; the mean, SD, and SE of the radius of gyration of the 60 NMR conformers are 11.03 Å, 0.06 Å, and 0.01 Å, respectively; the radius of gyration of the crystal structure is 10.70 Å. Extending these simulations from 316 ns^smt^ to 948 ns^smt^ yielded statistically the same results (Table III). The time series of radius of gyration for the GB3 conformers derived from the 20 948-ns^smt^ simulations using FF12MC or FF14SB do not show any signs of unfolding (Fig. S3). Clearly, both FF12MC and FF14SB were able to maintain the experimentally determined GB3 structure in the 20 948-ns^smt^ simulations.

**Table III.** Radii of gyration of experimental and simulated structures of folded globular proteins and related alpha carbon root mean square deviations between crystal structures and their corresponding NMR or simulated structures.

| Structure | Temp<br>(K) | No of<br>conformers | CαRMSD (Å) |  |  | RadGyr (Å) |  |  |
| --- | --- | --- | --- | --- | --- | --- | --- | --- |
|  |  |  | mean | SD | SE | mean | SD | SE |
| GB3 |  |  |  |  |  |  |  |  |
| 1IGD (X-ray) | ambient | 1 | — | — | — | 10.70 | — | — |
| 2LUM (NMR) | 298 | 1x60 | 0.80 | — | — | 11.03 | 0.06 | 0.01 |
| FF12MC | 297 | 20x1000 | 0.84 | 0.09 | 0.02 | 10.85 | 0.11 | 0.02 |
| FF14SBIm | 297 | 20x1000 | 0.89 | 0.09 | 0.02 | 10.97 | 0.11 | 0.02 |
| FF12MC | 297 | 20x3000 | 0.82 | 0.06 | 0.01 | 10.85 | 0.11 | 0.02 |
| FF14SBIm | 297 | 20x3000 | 0.86 | 0.05 | 0.01 | 10.97 | 0.11 | 0.02 |
| BPTI |  |  |  |  |  |  |  |  |
| 5PTI (X-ray) | ambient | 1 | — | — | — | 11.29 | — | — |
| 1PIT (NMR) | 309 | 1x20 | 1.18 | — | — | 11.37 | 0.07 | 0.02 |
| FF12MC | 309 | 20x1000 | 1.52 | 0.16 | 0.04 | 11.26 | 0.15 | 0.03 |
| FF14SBIm | 309 | 20x1000 | 0.89 | 0.15 | 0.03 | 11.48 | 0.10 | 0.02 |
| Ubiquitin |  |  |  |  |  |  |  |  |
| 1UBQ (X-ray) | ambient | 1 | — | — | — | 11.63 | — | — |
| 1D3Z (NMR) | 308 | 1x10 | 0.61 | — | — | 11.82 | 0.05 | 0.02 |
| FF12MC | 300 | 20x1000 | 1.54 | 0.32 | 0.07 | 11.66 | 0.15 | 0.03 |
| FF14SBIm | 300 | 20x1000 | 1.69 | 0.30 | 0.07 | 11.66 | 0.13 | 0.03 |
| FF12MC | 300 | 20x3000 | 1.69 | 0.31 | 0.07 | 11.68 | 0.20 | 0.04 |
| FF14SBIm | 300 | 20x3000 | 1.71 | 0.17 | 0.04 | 11.66 | 0.13 | 0.03 |
| Lysozyme |  |  |  |  |  |  |  |  |
| 4LZT (X-ray) | 295 | 1 | — | — | — | 14.03 | — | — |
| 1E8L (NMR) | 308 | 1x50 | 1.55 | — | — | 14.13 | 0.06 | 0.01 |
| FF12MC | 308 | 20x1000 | 1.7 | 0.7 | 0.2 | 14.21 | 0.28 | 0.06 |
| FF14SBIm | 308 | 20x1000 | 0.55 | 0.09 | 0.02 | 14.25 | 0.11 | 0.02 |

The GB3 simulations were repeated for ubiquitin using the same simulation conditions. These simulations were performed at 300 K that was used to spectroscopically determine the Lipari-Szabo order parameters of ubiquitin.^89^ The results of these simulations showed that FF12MC and FF14SB were able to maintain the experimentally determined ubiquitin structure in the 20 948-ns^smt^ simulations (Table III and Fig. S3). The GB3 simulations were also repeated for BPTI and lysozyme using the same simulation conditions. However, these simulations were not extended beyond 316 ns^smt^ because, unlike GB3 and ubiquitin, BPTI and lysozyme have multiple disulfide bonds restrain their folded conformations. These simulations were also performed at 309 K and 308 K, which were used in the experimental determination of the Lipari-Szabo order parameters of BPTI^91^ and lysozyme, respectively.^90^ The results also show that FF12MC and FF14SB are able to maintain the experimentally determined ubiquitin structure in the 20 316-ns^smt^ simulations (Table III and Fig. S3). Interestingly, according to CαRMSDs (Table III), the backbone conformations of BPTI and lysozyme in the FF14SB simulations are more restrained than those in the FF12MC simulations and those of the corresponding NMR structures. Taken together, the data in Table III and Fig. S3 show that, despite its weakness in reproducing main-chain *J*-coupling constants of GB3, ubiquitin, and lysozyme, FF12MC is able to simulate the experimentally determined conformations of GB3, BPTI, ubiquitin, and lysozyme in sub-microsecond NPT MD simulations.

### Simulating local motions of folded globular proteins

**Genuine localized disorders of BPTI and its mutant.** To investigate the ability of FF12MC and FF14SB to simulate the experimentally observed localized structural variations, the BPTI simulations were analyzed in the context of the report that the C14-C38 disulfide bond of BPTI adopts both left-and right-handed configurations^112^ in the NMR structure of PDB ID of 1PIT at 309 K^61^ (Fig. 5). Although C14-C38 of BPTI adopts the right-handed configuration in three different crystal structures (PDB IDs of 4PTI, 5PTI, and 6PTI),^61^ the C14-C38 flipping observed in the NMR study was confirmed later by a crystal structure of a BPTI mutant at the data-collection temperature of 290 K (PDB ID: 1QLQ).^66^ In this crystal structure, C14-C38 has the left-handed configuration with an occupancy parameter of 0.38 and the right-handed one with an occupancy parameter of 0.62. Further, the co-existence of two configurations for C14-C38 was also observed in an ultrahigh-resolution (0.86 Å) crystal structure of the same mutant at the data-collection temperature of 100 K (PDB ID: 1G6X).^113^

**Fig. 5.**
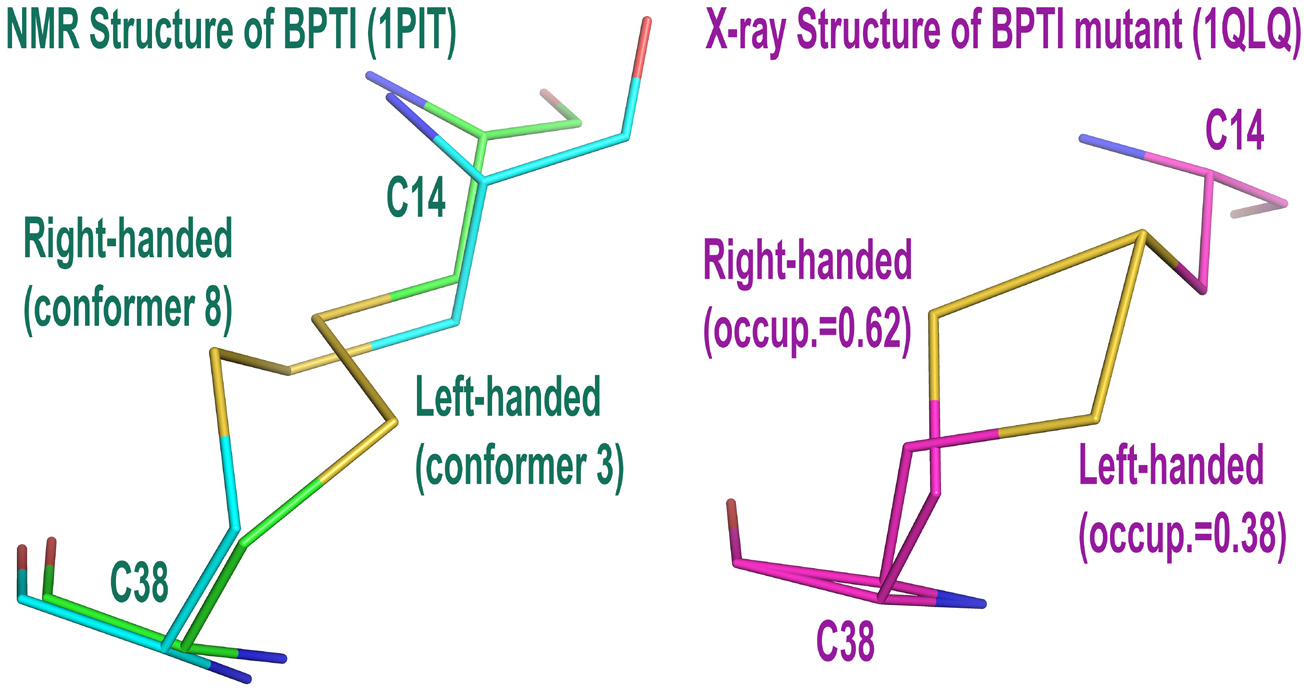
The right-and left-handed configurations of C14-C38 observed in the NMR structure of BPTI and the crystal structure of a BPTI mutant. The PDB IDs of the NMR and crystal structures are 1PIT and 1QLQ, respectively.

For the 20 simulations of BPTI using FF12MC at 309 K with the initial conformation taken from a 1PIT conformer that adopts the right-handed C14-C38 configuration, torsion cluster analysis showed that the most and the second most popular C14-C38 configurations over the first duration of 3.16 ns^smt^ were right-handed (occurrence of 38%) and left-handed (occurrence of 30%), respectively (Table S12). This trend remained when the analysis was repeated with durations extended to 31.6 ns^smt^ and 316 ns^smt^ (Table S12). Repeating the BPTI simulations at 290 K using the initial conformation taken from a 1QLQ conformer that adopts the left-handed C14-C38 configuration also showed the top-two most popular C14-C38 configurations to be right-handed (occurrence of 22%) and left-handed (occurrence of 16%) over the first 3.16-ns^smt^ duration. Extending these simulations to 31.6-ns^smt^ and 316-ns^smt^ yielded the same results except that the left-handed one became most popular during the two longer durations (Table S12).

For FF14SB, the 1PIT simulations resulted in the right-handed configuration being the sole configuration over the durations of 3.16 and 31.6 ns^smt^ and showed a mix of the right-handed configuration with an occurrence of 97% and the left one with an occurrence of 2% over the duration of 316 ns^smt^. The 20 simulations of 1QLQ using FF14SB under the same conditions as those for FF12MC showed that the right-handed configuration was absent over the duration of 3.16-ns^smt^ and present with occurrences of 1% and 2% over the durations of 31.6 and 316 ns^smt^, respectively (Table S12). The results suggest that FF14SB has the ability to lock C14-C38 into the right-handed configuration that was observed in the crystal structures of 4PTI, 5PTI, and 6PTI. These results also suggest that FF12MC has the ability to simulate the experimentally observed flipping between left-and right-handed configurations for C14-C38 of BPTI and its mutant, presumably due to the removal of torsions involving a nonperipheral *sp*^3^ atom. These unique abilities prompted the following studies to further compare the ability of the two forcefields to simulate subtle localized structural variations.

**The Lipari-Szabo order parameters.** The squared generalized order parameter (*viz*., the Lipari-Szabo order parameters) of a protein can be interpreted as a measure of the spatial restriction of an N-H bond in the protein, with the order parameter being 0 indicating the highest degrees of motion and 1 implying no motion.^65^ The main theorem of the order parameter is that two stochastic processes of global and local motions are separable by at least an order of magnitude; the global motions such as the overall tumbling correlation time (τ_c_) of a folded globular protein are on the timescale between a few ns^smt^ and tens of ns^smt^, whereas the local motions such as the motions of backbone N-H bonds are on the order of tens or hundreds of ps^smt^.^114^ In the context of this timescale of local motions, multiple sets of 20 standard-mass simulations that last up to 100 ns^smt^ using FF12MCsm and FF14SB were performed for calculation of the Lipari-Szabo order parameters of main-chain N-H bonds extracted from ^15^N spin relaxation data (S^2^) of GB3,^88^ ubiquitin,^89^ lysozyme,^90^ and BPTI^91^ to compare the ability of the two forcefields to simulate subtle backbone motions of folded globular proteins. In this study, Δ*t* = 1 fs^smt^ was used for simulations that lasted for 100 ns^smt^, while Δ*t* = 0.1 fs^smt^ was used for simulations that lasted for 50-500 ps^smt^. The reason to use FF12MCsm and Δ*t* = 0.1 fs^smt^ was to ensure adequate sampling in a short simulation.

According to RMSDs between computed and experimental S^2^ parameters (Table S13), FF12MCsm reproduced the experimental parameters of all four proteins with RMSDs ± SEs ranging from 0.063±0.005 to 0.074±0.002 on the timescale of 50 ps^smt^. For FFnMCsm, the S^2^ RMSDs of GB3 are insensitive to simulation time (Table S13). However, the S^2^ RMSDs of the other proteins do progress in time, and FFnMCsm best reproduced the experimental parameters of those proteins on the timescale of 50 ps^smt^ (Table S13). All S^2^ parameters calculated on the timescale of 50 ps^smt^ are shown in Fig. 3 with their standard errors listed in Table S4. By comparison, FF14SB reproduced the experimental parameters on the timescale of 50 ps^smt^ with RMSDs ± SEs ranging from 0.050±0.002 to 0.074±0.002, but it best reproduced the experimental data with RMSDs ± SEs ranging from 0.041±0.003 to 0.061±0.002 on the timescale of 4 ns^smt^ (Table S13). The S^2^ RMSDs of FF14SB are generally less sensitive to simulation time than those of FF12MCsm (Table S13). Although the S^2^ simulations using FF12MCsm and FF14SB were performed for up to 100 ns^smt^, the best calculated S^2^ parameters using FF12MCsm and FF14SB were not obtained on timescales that are close to five times the τ_c_s of the four proteins. This is partly because the stiffness of a protein exhibiting in the simulations using FFnMCsm or FF14SB differs from the stiffness using a forcefield — ff99SB_9φψ(g24;CS) −that led to the five times τ_c_ recommendation for best S^2^ estimation.^115^

Because the experimental S^2^ parameters were extracted from the ^15^N spin relaxation data on the picosecond timescale and the premise that the global and local motions are separable by at least an order of magnitude, the results of the nanosecond simulations suggest that FF12MC is on par with FF14SB in reproducing the experimental S^2^ parameters of GB3, ubiquitin, lysozyme, and BPTI on the timescale of 50 ps^smt^ (Fig. 3), although FF14SB better reproduces the experimental values than FF12MC on the timescale of 4 ns^smt^ that is in the range of the τ_c_s (2.0-5.7 ns^smt^) of the four proteins^88,90,116,117^ (Table S13). These results also prompted the following confirmation study on crystallographic B-factors that are akin to the S^2^ parameters.

**Crystallographic B-factors.** As a measure of the uncertainty of the atomic mean position, the crystallographic B-factor of a given atom reflects the displacement of the atom from its mean position in a crystal structure and this displacement attenuates X-ray scattering and is caused by both thermal motion of the atom and static disorder of the atom in a crystal lattice.^64,118–122^ Despite the challenges of separating the component of the thermal motion in time from the component of the disorder in space,^123^ crystallographic B-factors can often be used to quantitatively identify *less mobile* regions of crystal structures as long as the structures were determined without substantial crystal lattice defects, rigid-body motions, and refinement errors.^124,125^ A low B-factor indicates a small degree of motion, while a high B-factor may imply a large degree of motion.

In this context, to further evaluate the ability of FF12MC to simulate subtle thermal motions of a crystalline protein relative to that of FF14SB, simulated B-factors were obtained from atomic positional fluctuations that were calculated from 20 simulations of a folded globular protein in its solution state on the picosecond scale using FF12MCsm or FF14SB. Although simulations of proteins in their crystalline states^126,127^ can offer better and direct comparisons to the experimental data, simulations of proteins in the solution state were done in this study because the crystalline-state simulations are more computationally demanding than the solution-state simulations due to the larger size and slower convergence^127^ of the crystalline system. Further, in a reported study FF14SB was the best at reproducing experimental structural and dynamics properties among all four contemporary forcefields of FF99SB, FF14SB, FF14ipq, and CHARMM26.^127^ Direct comparison of FF12MC with FF14SB for their performances in the solution-state simulations can offer an insight into the ability of FF12MC to reproduce crystallographic B-factors.

Accordingly, the simulations for the S^2^ calculations were repeated at different temperatures. For GB3, BPTI, and ubiquitin, all simulations were performed at ambient temperature of 297 K because the exact data-collection temperatures of these proteins had not been reported. The lysozyme simulations were done at the reported data-collection temperature of 295 K.^128^

According to RMSDs between computed and experimental B-factors (Table S14), on the timescale of 50 ps^smt^, FF12MCsm best reproduces the experimental Cα and Cγ B-factors of all four proteins with RMSDs ± SEs ranging from 3.2±0.2 to 8±1 Å^2^ (average RMSD ± SE of 5.1±0.3 Å^2^) and from 7.8±0.8 to 9.9±0.7 Å^2^ (average RMSD ± SE of 9±2 Å^2^), respectively. On the timescale of 50 ps^smt^, FF14SB also best reproduces the experimental Cα and Cγ B-factors of all four proteins with RMSDs ± SEs ranging from 3.7±0.1 to 9±1 Å^2^ (average RMSD ± SE of 6.2±0.3 Å^2^) and from 8.5±0.3 to 10.3±0.2 Å^2^ (average RMSD ± SE of 9.1±0.5 Å^2^), respectively. For both FF12MC and FF14SB, the B-factor RMSDs of the BPTI and ubiquitin progress more in time than those of GB3 and lysozyme (Table S14). All Cα B-factors calculated on the timescale of on the timescale of 50 ps^smt^ are shown in Fig. 4 with their standard errors listed in Table S5.

These results show that FF12MC is on par with FF14SB in reproducing the crystallographic B-factors of the four proteins (Fig. 4). The results also demonstrate that FF12MC and FF14SB best reproduce the crystallographic B-factors on the timescale of 50 ps^smt^. This timescale corroborates the finding that FF12MC best reproduces the experimental S^2^ parameters on the timescale of 50 ps^smt^, suggesting that the calculated S^2^ parameters and B-factors on the 50-ps^smt^ timescale from the simulations using FF12MC and FF14SB may capture the true thermal fluctuations of folded globular proteins.

### How well FF12MC is trained to fold AAQAA

Given the encouraging results of FF12MC relative to those of FF14SB in all the afore-described studies except the main-chain protein *J*-coupling constant study, this FF12MC evaluation study turned to examining the ability of FF12MC to autonomously fold a short helical peptide AAQAA relative to those of FF12SB and FF14SB. Because AAQAA has been widely used for folding research and forcefield refinement,^12,13,84,129–140^ multiple sets of 20 simulations using FF12MC, FF12SB, and FF14SB to fold AAQAA were first carried out to determine how well FF12MC is trained—with a simple adjustment of two backbone scaling factors—to fold AAQAA relative to FF12SB and FF14SB.

Before describing the folding result of AAQAA, it is worth noting that AAQAA is not a typical α-helix peptide that exists in a mix of full α-helix, full coil, and central helices with frayed ends for at least three reasons. First, a small percentage of AAQAA was found to intermittently adopt conformations with an α-helix component at the *N*-terminus and a π-helix component in a region near the C-terminus,^129^ wherein the π helix is the 4.4_16_-helix found in 15% of known protein structures.^141^ Second, a study using a polarizable forcefield revealed a cooperative folding process in which the helical conformation is propagated throughout AAQAA once it is nucleated.^140^ Third, the NMR-derived residue distribution of helical content of AAQAA^51^ could not be predicted by the traditional Lifson-Roig model,^142–144^ a statistical mechanical model for theoretical prediction of the mean helical content (*viz*., fractional helicity) of a typical α-helix peptide. Side-chain•side-chain and side-chain•main-chain interactions had to be included in the traditional Lifson-Roig model to correctly simulate the experimentally observed residue distribution of helical content of AAQAA.^51^

Therefore, no attempt was made to compute the Lifson-Roig parameters of AAQAA in the present study. As described in METHODS, the fractional helicity of AAQAA was estimated from torsions *ϕ* and *ψ* using a simple protocol that is based on local α-helix content of four consecutive residues. Because of a considerable overlap of the *ϕ* and *ψ* torsions between α- and π-helices^76^ and the subjective nature of defining the torsion ranges of *ϕ* and *ψ* for identification of 3_10_-, α-, and π-helices, no attempt was made to include all components of the three helices in the fractional helicity estimation.

To substantiate the torsion-based protocol, the mean helix content was also estimated from the global α-helix content of AAQAA according to CαβRMSD from the most populated helical conformation (see METHODS). This alternative is reasonable as long as the population of the second most populated conformation of AAQAA is substantially smaller than the population of the most populated helical conformation. If the fractional helicity derived from the first protocol is slightly higher than the one derived from the second protocol, both protocols are considered to be reasonable. Indeed, according to the cluster analysis of 20 3.16-μs^smt^ simulations of AAQAA using FF12MC at 274 K (Table S3), the representative, instantaneous conformation in the largest cluster of AAQAA at 274 K is a full-α-helix conformation (Fig. 2A) with a population of 41.7% (Table S3). The second most populated conformation at 274 K has Ala1-Ala5 adopting an α-helix, Ala2-Ala7 adopting a hybrid between α-helix and π helix, Ala3-Ala9 adopting a π helix, and Ala6-Ala15 adopting a π helix (Fig. 2B). This conformation has a population of 8.0% (Table S3). The third most popular conformation at 274 K is a partial-α-helix conformation with frayed residues of Ac, Ala1, and NH_2_ (Fig. 2C), and this conformation has a population of 3.5% (Table S3). The populations of the three most popular conformations with the full-, mixed-, and partial-α-helix conformation decreased to 18.9%, 5.8%, and 3.0% at 300 K and 14.4%, 5.6%, and 2.1% at 310 K, respectively (Table S3), but the rank orders of these populations at 300 K and 310 K are the same as the one at 274 K. These results support the use of the alternative protocol to estimate the mean helix content of AAQAA and the use of the most popular full-α-helix conformation (Fig. 2A) as the native conformation of AAQAA for the autonomous folding study described below.

According to the smoothed time series of CαβRMSD from the native conformation, the aggregated native state populations, and the estimated folding times of AAQAA at different temperatures (Table IV and Fig. S1A-C), FF12MC, FF12SB, and FF14SB can autonomously fold AAQAA from a fully extended backbone conformation to the native conformation and simulate subsequent unfolding and refolding in all (for FF12MC) or some (for FF12SB and FF14SB) of 20 simulations at 274 K.

**Table IV.** Folding of hairpin and helical peptides and a miniprotein Trp-cage in isothermal-isobaric molecular dynamics simulations at 1 atm.

| Sequence | Temperature<br>(K) | Aggregated<br>simulation<br>time ($\mu\text{s}^{\text{smt}}$) | Aggregated native<br>state population (%) | | | Estimated folding time ($\text{ns}^{\text{smt}}$) | | | |
| --- | --- | --- | --- | --- | --- | --- | --- | --- | --- |
|  |  |  | mean | SD | SE | mean | LCL | UCL | Event |
| FF12MC |  |  |  |  |  |  |  |  |  |
| Chignolin | 277 | 20 x 3.16 | 47 | 11 | 2 | 153 | 99 | 237 | 20 |
| Chignolin | 300 | 20 x 3.16 | 33 | 10 | 2 | 79 | 51 | 123 | 20 |
| CLN025 | 277 | 20 x 3.16 | 70 | 15 | 3 | 433 | 279 | 671 | 20 |
| CLN025 | 300 | 20 x 3.16 | 63 | 13 | 3 | 174 | 112 | 270 | 20 |
| AAQAA | 274 | 20 x 3.16 | 41 | 8 | 2 | 189 | 122 | 293 | 20 |
| AAQAA | 300 | 20 x 3.16 | 18 | 3 | 1 | 143 | 92 | 221 | 20 |
| AAQAA | 310 | 20 x 3.16 | 14 | 3 | 1 | 92 | 59 | 142 | 20 |
| Trp-cage | 280 | 30 x 8.848 | 18 | 8 | 1 | 1998 | 1396 | 2860 | 30 |
| Chignolin | 277 | 20 x 1.00 | 40 | 18 | 4 | 153 | 99 | 237 | 20 |
| CLN025 | 277 | 20 x 1.00 | 41 | 31 | 7 | 446 | 281 | 708 | 18 |
| AAQAA | 274 | 20 x 1.00 | 38 | 13 | 3 | 189 | 122 | 293 | 20 |
| FF12SB |  |  |  |  |  |  |  |  |  |
| Chignolin | 277 | 20 x 1.00 | 3 | 7 | 2 | 871 | 506 | 1500 | 13 |
| CLN025 | 277 | 20 x 1.00 | 4 | 10 | 2 | – | – | – | 4 |
| AAQAA | 274 | 20 x 1.00 | 2 | 7 | 2 | 1287 | 712 | 2326 | 11 |
| AAQAA | 300 | 20 x 1.00 | 5 | 6 | 1 | 416 | 265 | 651 | 19 |
| AAQAA | 310 | 20 x 1.00 | 4 | 3 | 1 | 250 | 159 | 391 | 19 |
| FF12SBIm |  |  |  |  |  |  |  |  |  |
| CLN025 | 277 | 20 x 3.16 | 22 | 29 | 6 | 3328 | 1889 | 5863 | 12 |
| FF14SB |  |  |  |  |  |  |  |  |  |
| Chignolin | 277 | 20 x 1.00 | 19 | 30 | 7 | 550 | 342 | 886 | 17 |
| CLN025 | 277 | 20 x 1.00 | 7 | 14 | 3 | 1012 | 600 | 1708 | 14 |
| AAQAA | 274 | 20 x 1.00 | 4 | 5 | 1 | 1224 | 677 | 2213 | 11 |
| FF14SBIm |  |  |  |  |  |  |  |  |  |
| CLN025 | 277 | 20 x 3.16 | 38 | 30 | 7 | 1366 | 860 | 2170 | 18 |

SD: Standard deviation. SE: Standard error. LCL: Lower 95% confidence limit. UCL: Upper 95% confidence limit. <sup>smt</sup>: Standard-mass time. Event: The number of simulations that captured a folding event.

For the 20 3.16-μs^smt^ low-mass simulations of AAQAA using FF12MC at Δ*t* = 1.00 fs^smt^, the aggregated native state populations (*viz*., the α-helix populations) and their SDs are 41±8% at 274 K, 18±3% at 300 K, and 14±3% at 310 K (Table IV). These populations are slightly smaller than the estimated fractional helicities of 55±6% at 274 K, 35±3% at 300 K, and 29±4% at 310 K (Table V). Both the α-helix populations and the estimated fraction helicities are in reasonable agreement with the experimentally observed fractional helicities ± SDs of 50.6±0.4% at 274 K, 20.8±0.4% at 300 K, and 13.5±0.4% at 310 K (Table V). The τ_f_s of AAQAA predicted from the 20 simulations using FF12MC are 189 ns^smt^ (95% CI: 122-293 ns^smt^) at 274 K, 143 ns^smt^ (95% CI: 92221 ns^smt^) at 300 K, and 92 ns^smt^ (95% CI: 59-142 ns^smt^) at 310 K, respectively (Table IV). These small SDs and narrow 95% CIs of the 20 simulations relative to the means suggest that the simulations using FF12MC are converged. This convergence is also supported by the smoothed time series of CαβRMSD (Fig. S1A) showing that all simulations captured the most popular full-α-helix conformation.

**Table V.**
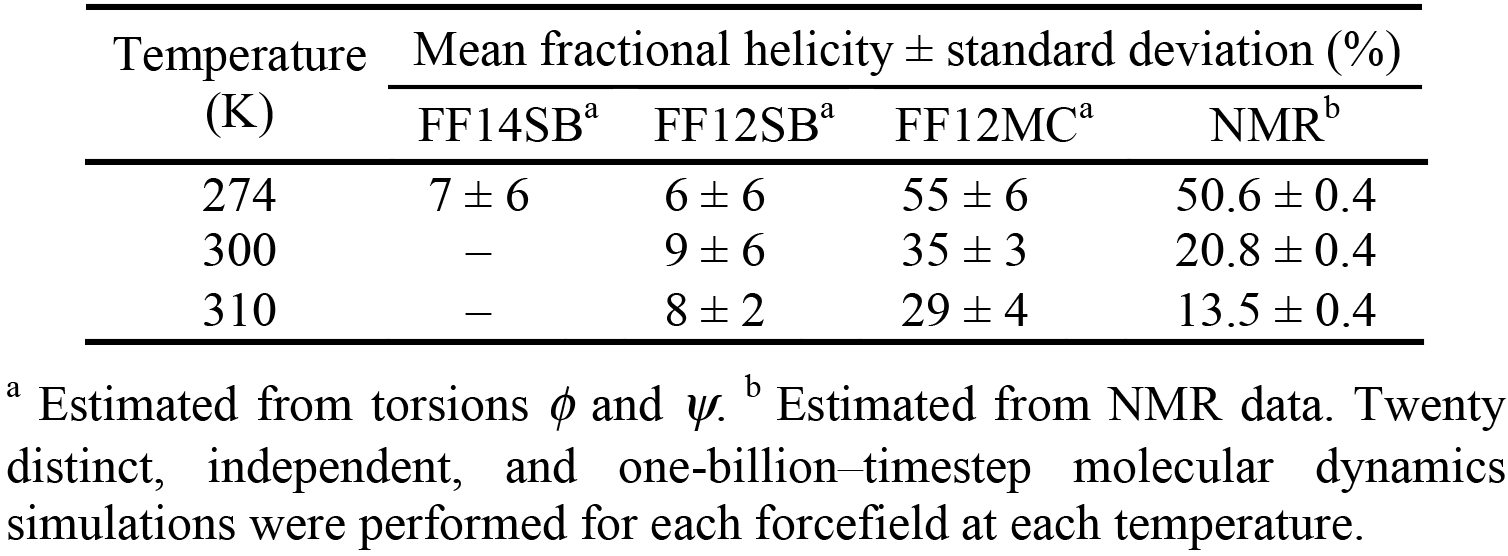
Mean fractional helicity of Ac-(AAQAA)_3_-NH_2_ estimated from NMR data and MD simulations.

| Temperature<br>(K) | Mean fractional helicity ± standard deviation (%) | | | |
| --- | --- | --- | --- | --- |
|  | FF14SB<sup>a</sup> | FF12SB<sup>a</sup> | FF12MC<sup>a</sup> | NMR<sup>b</sup> |
| 274 | 7 ± 6 | 6 ± 6 | 55 ± 6 | 50.6 ± 0.4 |
| 300 | – | 9 ± 6 | 35 ± 3 | 20.8 ± 0.4 |
| 310 | – | 8 ± 2 | 29 ± 4 | 13.5 ± 0.4 |

<sup>a</sup> Estimated from torsions $\phi$ and $\psi$ . <sup>b</sup> Estimated from NMR data. Twenty distinct, independent, and one-billion-timestep molecular dynamics simulations were performed for each forcefield at each temperature.

For the 20 1.00-μs^smt^ standard-mass simulations of AAQAA using FF12SB at Δ*t* = 1.00 fs^smt^, the aggregated native state populations ± SDs are 2±7% at 274 K, 5±6% at 300 K, and 4±3% at 310 K (Table IV). The corresponding population ± SD of FF14SB is 4±5% at 274 K (Table IV). Very low fractional helicities of AAQAA were also observed in the simulations using FF12SB and FF14SB (Table V). The τ_f_s of AAQAA predicted from the 20 simulations using FF12SB are 1287 ns^smt^ (95% CI: 712-2326 ns^smt^) at 274 K, 416 ns^smt^ (95% CI: 265-651 ns^smt^) at 300 K, and 250 ns^smt^ (95% CI: 159-391 ns^smt^) at 310 K, respectively (Table IV). The τ_f_ of AAQAA predicted from the 20 simulations using FF14SB is 1224 ns^smt^ (95% CI: 677-2213 ns^smt^) at 274 K (Table IV). These large SDs and wide 95% CIs relative to the means show that an aggregated simulation time of 20 μs^smt^ is inadequate to estimate the native state populations, mean helix content, or folding times of FF12SB and FF14SB. This conclusion is consistent with the smoothed time series of CαβRMSD showing that only some of 20 simulations captured the full-α-helix conformation (Table IV and Fig. S1B and C).

As explained above, FF12MC was used in the low-mass simulations at Δ*t* = 1.00 fs^smt^ (viz., Δ*t* = 3.16 fs^lmt^), whereas FF12SB and FF14SB were used in the standard-mass simulation at Δ*t* = 1.00 fs^smt^. This was so that FF12SB and FF14SB were evaluated with higher integration accuracy than the accuracy used for FF12MC. Therefore, the results of the 20 1.00-μs^smt^ standard-mass simulations using FF12SB or FF14SB should be compared with those of 20 1.00-μs^smt^ low-mass simulations using FF12MC. As listed in Table IV, the aggregated native state population ± SD (38±13%) of 20 1-μs^smt^ low-mass simulations of AAQAA at 274 K using FF12MC is significantly higher than that (2±7% or 4±5%) of the 20 1-μs^smt^ standard-mass simulations using FF12SB or FF14SB, respectively (Table IV). The τ_f_ (189 ns^smt^) of 20 1-μs^smt^ low-mass simulations at 274 K using FF12MC is also significantly shorter than that (1287 ns^smt^ or 1224 ns^smt^) of the 20 1-μs^smt^ standard-mass simulations using FF12SB or FF14SB. The two sets of simulations with equal aggregated simulation times show that FF12SB and FF14SB cannot fold AAQAA as fast as FF12MC. This is hardly surprising because FF12SB and FF14SB were not benchmarked against AAQAA,^16^ but it shows that by a simple adjustment of two backbone scaling factors FF12MC is well trained to autonomously fold AAQAA.

### Folding, unfolding, and refolding of β-hairpins

FF12MC, FF12SB, and FF14SB were then tested for their ability to autonomously fold β-hairpins of chignolin and CLN025. According to the smoothed time series of CαβRMSD from the native conformations, the aggregated native state populations, and the estimated folding times (Table IV and Fig. S1D-F), all three forcefields can fold the two β-hairpins from fully extended backbone conformations to their native conformations and simulate subsequent unfolding or partial unfolding and refolding in all (for FF12MC) or some (for FF12SB and FF14SB) of 20 1.00-μs^smt^ simulations at Δ*t* = 1.00 fs^smt^ and 277 K. Cluster analysis shows that the average chignolin conformation of the largest cluster identified from the simulations using FF12MC, FF12SB, or FF14SB has a CRMSD from the first model of the NMR structure^67^ of 1.62 Å, 3.32 Å, or 1.33 Å, respectively (Fig. 1E, C, *and* D). The CRMSD of the average chignolin conformation of the second largest cluster of FF12SB was 1.54 Å (Table S3). When compared to the NMR structure of CLN025,^68^ the average CLN025 conformation of the largest cluster of the simulations using FF12MC, FF12SB, or FF14SB has a CRMSD of 1.72 Å, 1.76 Å, or 1.70 Å, respectively (Fig. 1J, H, and I). When compared to the crystal structure of CLN025,^68^ the CRMSDs increased to 2.68 Å, 2.59 Å, and 2.51 Å, respectively. These CRMSDs show that all three forcefields can fold CLN025 in water to conformations that resemble more the solution structure than the crystalline structure.

Despite the removal of torsions that involve a nonperipheral *sp^3^* atom in FF12MC and the use of At = 3.16 fs^lmt^ for FF12MC and Δ*t* = 1.00 fs^smt^ for FF14SB, the CRMSD between two average CLN025 conformations of the most populated clusters derived from the simulations using FF14SB and FF12MC was 0.45 Å, whereas the CRMSD between the NMR and crystal structures of CLN025 was 3.15 Å. In addition, the aggregated native state populations ± SDs of chignolin and CLN025 in the 20 3.16-μs^smt^ low-mass simulations at 277 K using FF12MC were 47±11% and 70±15%, respectively (Table IV). The corresponding populations ± SDs reduced to 33±10% and 63±13%, respectively, when the simulation temperature increased to 300 K (Table IV). These significant differences are consistent with the experimental study showing that CLN025 is more stable than the parent protein chignolin.^68^ The small SDs of the 20 simulations relative to the means of the native state populations suggest that the simulations using FF12MC are converged. The convergence is also supported by the smoothed time series of CαβRMSD (Fig. S1D) showing that all 20 simulations using FF12MC captured the folding of chignolin or CLN025. The τ_f_s of chignolin and CLN025 predicted from the 20 simulations at 277 K using FF12MC were 153 ns^smt^ (95% CI: 99-237 ns^smt^) and 433 ns^smt^ (95% CI: 279-671 ns^smt^), respectively (Table IV). Furthermore, the τ_f_ of CLN025 estimated from the 20 simulations using FF12MC was 174 ns^smt^ (95% CI of 112-270 ns^smt^) at 300 K (Table IV). This τ_f_ −obtained by using the Kaplan-Meier estimator without any prior knowledge of the hazard function for the nonnative state population of CLN025—agrees with the experimental study showing that CLN025 folds with a τ_f_ of approximately 100 ns^smt^.^7^° This agreement suggests that FF12MC may have adequately sampled nonnative states of CLN025 in the 20 simulations at 300 K, which is consistent with the unique ability of FF12MC to simulate the genuine disorder of C14-C38 in BPTI and its mutant.

The aggregated native state populations ± SDs of chignolin and CLN025 in the 20 1.00-μs^smt^ simulations at Δ*t* = 1.00 fs^smt^ and 277 K using FF12SB (or FF14SB) were 3±7% (or 19±30%) and 4±10% (or 7±14%), respectively (Table IV). The τ_f_ of chignolin predicted from the 20 simulations using FF12SB was 871 ns^smt^ (95% CI: 506-1500 ns^smt^) at 277 K (Table IV). The τ_f_ of CLN025 at 277 K for FF12SB could not be estimated with confidence because more than half of the 20 simulations did not capture a folding event. The τ_f_s of chignolin and CLN025 predicted from the 20 simulations using FF14SB at 277 K were 550 ns^smt^ (95% CI: 342-886 ns^smt^) and 1012 ns^smt^ (95% CI: 600-1708 ns^smt^), respectively (Table IV). These large SDs and wide 95% CIs relative to the means indicate poor convergence of the simulations using FF12SB and FF14SB. This poor convergence is consistent with the number of simulations that captured a folding event listed in Table IV and the smoothed time series of CαβRMSD in Fig. S1E and F showing that some simulations did not capture the native conformations.

As listed in Table IV, the aggregated native state populations of chignolin and CLN025 ± SEs obtained from 20 1.00-μs^smt^ low-mass simulations at 277 K using FF12MC at Δ*t* = 1.00 fs^smt^ were 40±4% and 41±7%, respectively; the τ_f_s of chignolin and CLN025 estimated from the 20 1.00-μs^smt^ low-mass simulations at 277 K using FF12MC were 153 ns^smt^ (95% CI: 99-237 ns^smt^) and 446 ns^smt^ (95% CI: 281-708 ns^smt^), respectively. By comparison, the aggregated native state populations of chignolin and CLN025 ± SEs obtained from 20 1.00-μs^smt^ standard-mass simulations at 277 K using FF12SB (FF14SB) at Δ*t* = 1.00 fs^smt^ were 3±2% (19±7%) and 4±2% (7±3%), respectively; the τ_f_s of chignolin estimated from the 20 1.00-μs^smt^ standard-mass simulations at 277 K using FF12SB and FF14SB were 871 ns ^smt^ (95% CI: 506-1500 ns^smt^) and 550 ns ^smt^ (95% CI: 342-886 ns^smt^), respectively; the corresponding τ_f_ of CLN025 for FF14SB was 1012 ns ^smt^ (95% CI: 600-1708 ns^smt^).

Further, 20 3.16-μs^smt^ low-mass simulations at Δ*t* = 1.00 fs^smt^ and 277 K were performed to fold CLN025 using FF12SBlm or FF14SBlm, wherein FF12SBlm and FF14SBlm denote the respective forcefields with their atomic masses reduced uniformly by tenfold. The resulting populations ± SEs of CLN025 (22±6% for FF12SBlm and 38±7% for FF14SBlm) derived from the 20 3.16-μs^smt^ low-mass simulations are still significantly lower than the corresponding one (70±3%) for FF12MC (Table IV). The resulting τ_f_s of CLN025 at 277 K (3328 ns^smt^ and 95% CI: 1889-5863 ns^smt^ for FF12SBlm; 1366 ns^smt^ and 95% CI: 860-2170 ns^smt^ for FF14SBlm) are also substantially longer than that (433 ns^smt^ and 95% CI: 279-671 ns^smt^) estimated from the 20 3.16- μs^smt^ low-mass simulations using FF12MC (Table IV). These results show that FF12MC can indeed fold the two β-hairpins with folding times that are both shorter than those using FF12SB and FF14SB and closer to the experimental values.

### Folding, unfolding, and refolding of an α-miniprotein

To evaluate the ability of FF12MC to fold an α-miniprotein that is larger than the β-hairpins, autonomous folding simulations of a 20-residue Trp-cage (the TC10b sequence^69^) were carried out at 280 K at which the NMR structure of TC10b was determined. FF12SB and FF14SB were not included in this computationally demanding study because as noted above these forcefields fold chignolin, CLN025, and AAQAA at much slower rates than FF12MC. According to the smoothed time series of CαβRMSD from the native conformation, the aggregated native state population, and the estimated folding time (Table IV and Fig. S1G), FF12MC can fold the Trp-cage from a fully extended backbone conformation to its native conformation and simulate subsequent unfolding and refolding in all 30 8.848-μs^smt^ low-mass simulations at Δ*t* = 1.00 fs^smt^ and 280 K, with (*i*) an aggregated native state population ± SD of 18±8% (Table IV), (*ii*) a τ_f_ of 1998 ns^smt^ (95% CI: 1396-2860 ns^smt^) at 280 K (Table IV), and (*iii*) a CRMSD of 1.53 Å between the first NMR model and the average conformation of the largest cluster identified from the Trp-cage simulations (Fig. 1O). More importantly, the τ_f_s of 1998 ns^smt^ at 280 K for the Trp-cage (TC10b) — obtained by using the Kaplan-Meier estimator without any prior knowledge of the hazard function for the nonnative state population of the miniprotein — is consistent with the experimentally determined τ_f_ of 1430 ns^smt^ at 300 K.^71^ Plotting the natural logarithm of the nonnative state population versus time-to-folding from nonnative states to the native state of the Trp-cage reveals a linear relationship with r^2^ of 0.9408 (Fig. 6), indicating an exponential decay of the nonnative state population of the Trp-cage over simulation time. This exponential decay is in excellent agreement with the experimental observation that the folding of Trp-cage follows a two-state kinetics scheme.^71^ These results show that FF12MC can fold the Trp-cage from scratch with high accuracy in the 30 simulations at 280 K. Further, the results demonstrate that FF12MC can capture the two-state kinetics scheme as the major folding pathway of the Trp-cage with an estimated τ_f_ that is consistent with the experimental value.

**Fig. 6.**
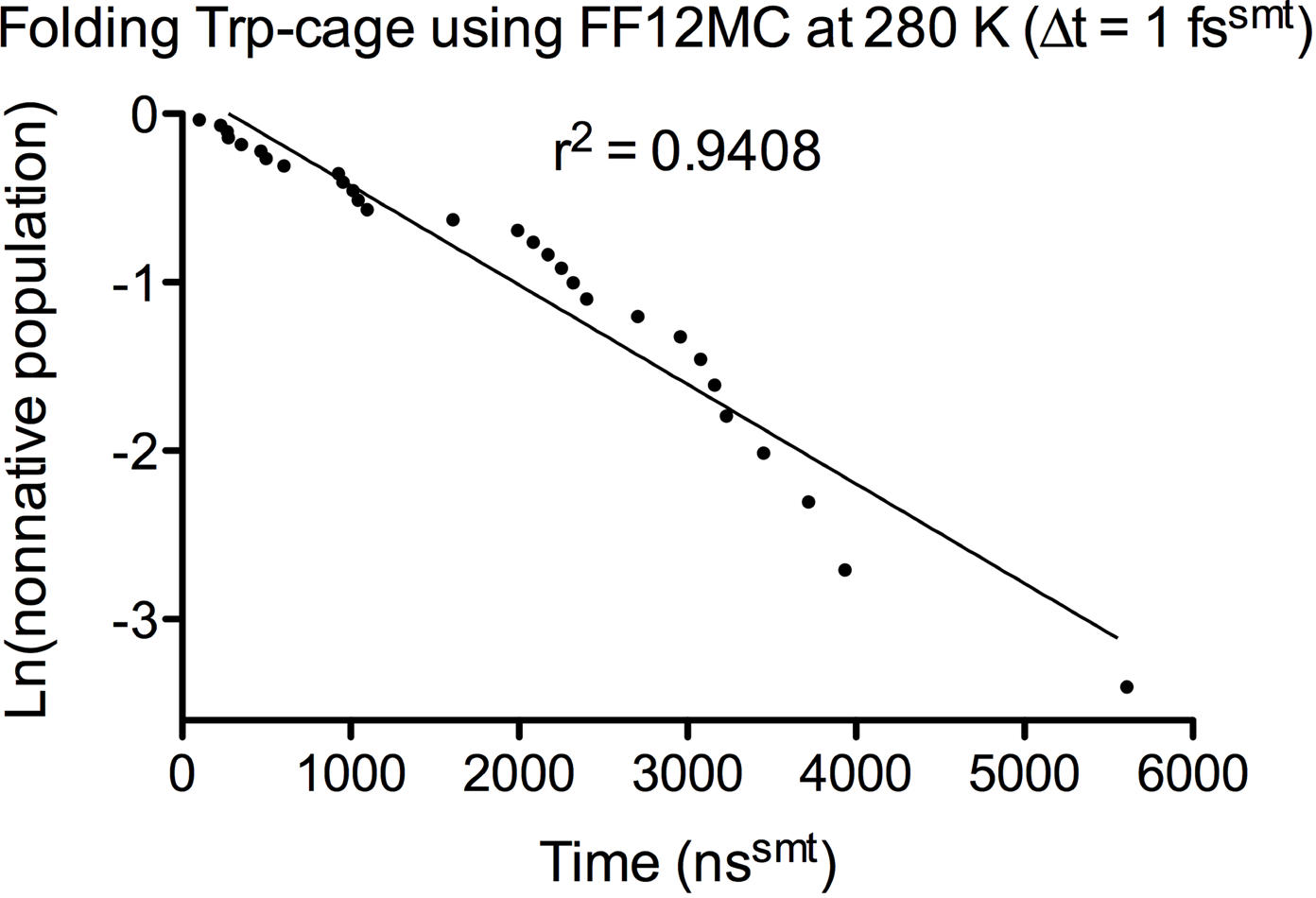
Plot of the natural logarithm of the nonnative state population of the Trp-cage (TC10b) over time-to-folding. The individual folding times were taken from the data provided in Fig. S1G. The linear regression analysis was performed using the PRISM 5 program.

### Refining CASPR models TMR01, TMR04, and TMR07

Consistent with the need to use statistically-derived knowledge-based potentials for refining comparative models of protein structure,^31,37,39,145–149^ the accuracy of the physics-based forcefield — such as those in the form of Eq. 1 — has been suggested to be the primary factor limiting the simulation-based comparative model refinement.^41^ This inspired a simulation-based refinement study of comparative models of monomeric globular proteins to compare FF12MC with FF14SB and FF96^57^ for their ability to generate conformations that cluster around the native conformation of a test protein. While better refinement can be achieved by performing restricted MD simulations,^28,32,41,44^ unrestricted and unbiased NPT MD simulations were performed in this study because of its objective to evaluate the effectiveness of a forcefield rather than a refinement protocol.

This refinement study used models TMR01, TMR04, and TMR07 of the first CASPR experiment. Four other models of the experiment were excluded for the following reasons. TMR02 and TMR03 are in the monomeric form, but their crystal structures are in multimeric forms (PDB IDs: 1VMO and 1VLA). A calcium ion is missing in TMR05 but present in the corresponding crystal structure (PDB ID: 1TVG). TMR06 has a VahMet mutation and deletion of residues from -8 to 0 relative to the corresponding crystal structure (PDB ID: 1XG8). The refinement studies of TMR01, TMR04, and TMR07 by replica-exchange MD simulations using a physics-based forcefield GBSW and a knowledge-based function RAPDF/HB_EM_ have been reported.^32,37^ These studies serve as valuable benchmarks for the present study.

FF96 was chosen because of the insight this early forcefield can offer into how much improvement the AMBER forcefield has made from FF96 to FF14SB and FF12MC over the past two decades. It was chosen also because the refinement of TMR01 made by this author using FF96 and the low-mass sampling enhancement technique earned a top score (ΔCαRMSD of −2.853 Å) in the first CASPR experiment in 2006 (see Supporting Information Note S1 for the CASPR organizers’ assessment). To justify its use, FF12MC must perform substantially better in refining TMR01 than FF96lm, wherein FF96lm denotes FF96 with its atomic masses reduced uniformly by tenfold.

In this refinement study, each refined model was assessed by quality scores (QSs) of GDT-HA,^95^ GDC-all,^96^ RPF,^97^ and LDDT.^98^ These QSs were used for the assessment of comparative model refinement of CASP10.^97^ However, the reported refinement studies^32,37^ of TMR01, TMR04, and TMR07 used sseRMSD, CαRMSD, and GDT-TS. To facilitate comparison, sseRMSD, CαRMSD, and GDT-TS were also included in the present study. While sseRMSD, CαRMSD, GDT-TS, GDT-HA, and GDC-all are five QSs based on global alignment, RPF and LDDT are two QSs based on local alignment. The SphereGrinder score^99^ and the CAD score^100^ were therefore included to balance the local-alignment scores with the global-alignment scores. Hereafter, RPF9 and LDDT15 denote the RPF and LDDT scores that were calculated with a distance cutoff of 9.0 Å and 15.0 Å, respectively; SG2n6 denotes the SphereGrinder score that was calculated with an all-atom RMSD cutoff of 2.0 Å and a sphere radius of 6.0 Å. The MolProbity score^150^ was excluded in the model assessment of CASP10 because a perfect α-helix prediction can have an excellent MolProbity score even though the experimental structure is a β-stand.^97^ Therefore, the MolProbity score was excluded in this study.

As shown in Fig. 7, relative to the crystal structure (PDB ID: 1XE1), the CαRMSD, GDT-HA, and SG2n6 scores of TMR01 are 6.1 Å, 0.593, and 0.495, respectively (Table VI). These QSs are mainly due to large conformational differences at the N-terminus (residues 18-25 of 1XE1) and three loops (residues 35-42 of 1XE1 for Loop 1; residues 59-64 of 1XE1 for Loop 2; residues 89-97 of 1XE1 for Loop 3). Refining TMR01 using 20 316-ns^smt^ low-mass simulations at 340 K and Δ*t* = 1.00 fs^smt^ using FF12MC substantially improved the CαRMSD, GDT-HA, and SG2n6 scores to 1.4 Å, 0.797, and 0.766, respectively (Table VI). The refinement using FF14SBlm and the same simulation conditions of FF12MC improved the CαRMSD, GDT-HA, and SG2n6 scores to 3.0 Å, 0.717, and 0.663, respectively. Under the same simulation conditions, FF96lm also considerably improved the CαRMSD, GDT-HA, and SG2n6 scores of TMR01 to 3.9 Å, 0. 712, and 0.629, respectively (Table VI). Using the same refinement protocol as the one for TMR01, all three forcefields considerably improved all nine QSs of TMR04 and TMR07 except that FF12MC and FF96lm slightly increased CαRMSD from 2.2 Å to 2.4 Å and 2.7 Å, respectively, for TMR07 (Fig. 7 and Table VI). An increase of CαRMSD to 2.7 Å was also observed in the TMR07 refinement by GBSW (Table VI). The performance differences among the three forcefields for TMR04 and TMR07 are not as large as those for TMR01. This is because the refinement of TMR01 involves a much larger conformational change than those involved in the refinement of TMR04 and TMR07, as indicated by the respective CαRMSDs of Table VI.

**Fig. 7.**
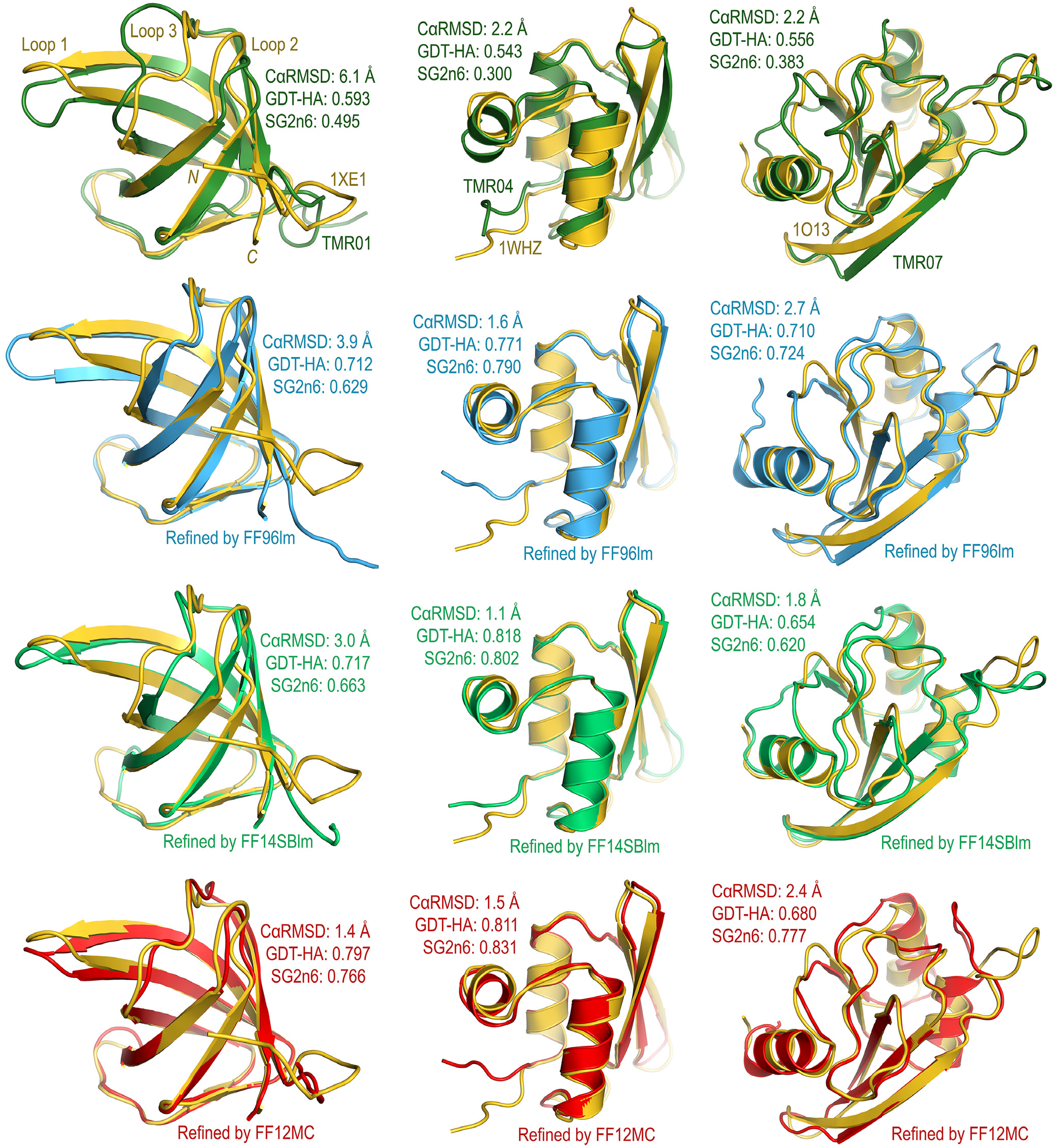
Overlays of three CASPR crystal structures with unrefined and refined models. The Protein Data Bank IDs of the crystal structures of TMR01, TMR04, and TMR07 are 1XE1, 1WHZ, and 1O13, respectively. Each refined model is the average conformation of the largest cluster of 20 unbiased, unrestricted, distinct, independent, and 316-ns^smt^ NPT MD simulations of a CASPR model at Δt = 1.00 fs^smt^ and 340 K using FF12MC, FF14SBlm, or FF96lm.

**Table VI.** Quality Scores for Refining Three CASPR Models by Five Different Forcefields.

| Model | Refinement | sseRMSD<br>(Å) | CαRMSD<br>(Å) | GDT-TS | GDT-HA | GDC-all | RPF9 | LDDT15 | SG2n6 | CAD |
| --- | --- | --- | --- | --- | --- | --- | --- | --- | --- | --- |
| TMR01 | None | 1.3 | 6.1 | 0.772 | 0.593 | 0.491 | 0.690 | 0.631 | 0.495 | 0.609 |
|  | RAPDF/HB<sub>EM</sub> | 0.9 | – | – | – | – | – | – | – | – |
|  | GBSW | – | 3.9 | 0.835 | – | – | – | – | – | – |
|  | FF12MC | 0.7 | 1.4 | 0.920 | 0.797 | 0.820 | 0.851 | 0.791 | 0.766 | 0.693 |
|  | FF14SBIm | 1.1 | 3.0 | 0.849 | 0.717 | 0.689 | 0.805 | 0.750 | 0.663 | 0.656 |
|  | FF96Im | 1.4 | 3.9 | 0.854 | 0.712 | 0.653 | 0.776 | 0.723 | 0.629 | 0.642 |
| TMR04 | None | 1.8 | 2.2 | 0.743 | 0.543 | 0.637 | 0.667 | 0.603 | 0.300 | 0.626 |
|  | RAPDF/HB<sub>EM</sub> | 0.8 | – | – | – | – | – | – | – | – |
|  | GBSW | – | 1.6 | 0.900 | – | – | – | – | – | – |
|  | FF12MC | 0.6 | 1.5 | 0.932 | 0.811 | 0.793 | 0.800 | 0.762 | 0.831 | 0.687 |
|  | FF14SBIm | 0.8 | 1.1 | 0.939 | 0.818 | 0.838 | 0.805 | 0.776 | 0.802 | 0.683 |
|  | FF96Im | 0.7 | 1.6 | 0.921 | 0.771 | 0.776 | 0.777 | 0.744 | 0.790 | 0.664 |
| TMR07 | None | 1.9 | 2.2 | 0.766 | 0.556 | 0.668 | 0.686 | 0.618 | 0.383 | 0.590 |
|  | RAPDF/HB<sub>EM</sub> | 2.1 | – | – | – | – | – | – | – | – |
|  | GBSW | – | 2.7 | 0.810 | – | – | – | – | – | – |
|  | FF12MC | 1.5 | 2.4 | 0.846 | 0.680 | 0.732 | 0.815 | 0.762 | 0.777 | 0.694 |
|  | FF14SBIm | 1.2 | 1.8 | 0.832 | 0.654 | 0.753 | 0.762 | 0.713 | 0.620 | 0.689 |
|  | FF96Im | 1.6 | 2.7 | 0.872 | 0.710 | 0.719 | 0.793 | 0.765 | 0.724 | 0.711 |

To rank the performances of FF12MC, FF14SBlm, FF96lm, RAPDF/HB_EM_, and GBSW in refining TMR01, TMR04, and TMR07, this study used two standardization protocols (classical and robust Z scores) that were used to rank the model refinement groups in CASP experiments 9 and 10.^97,151^ For each of the three CASPR models refined by M number of forcefields, a QS-specific Z score was calculated for each of N number of QSs. Averaging all N QS-specific Z scores of each model with an equal weight gave rise to a model-specific Z score. Averaging all three model-specific Z scores of each forcefield with an equal weight gave rise to a forcefield-specific Z score (Z^F^). The classical SD-based Z score was calculated according to Eq. 2. To minimize the influence of “outliers,” the robust Z score that is based on median absolute deviation about the median^152^ was calculated according to Eq. 3, wherein med(QS_*i*,M_) is the median of {QS_*i*,1_,QS_*i*,2_, … QS_*i,j*_, …, QS_*i*,M_} and *i* ∈ {1, 2, …, N}, QS_*i,j*_ is the QS_*i*_ of forcefield *j*, and med(|QS_*i*,M_ - med(QS_*i*,M_)|) is the median of {|QS_*i*,1_ - med(QS_*i*,M_)|, QS_*i*,2_ - med (QS_*i*,M_)|,…, |QS_*i,j*_ - med(QS_*i*,M_)|, …, |QS_*i*,M_ - med(QS_*i*,M_)|}. Missing QSs were assigned a QS-specific Z score of zero, in the same way as it was done for the model assessment of CASP10.^97^

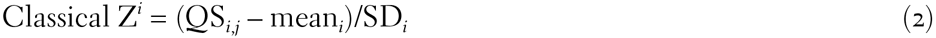

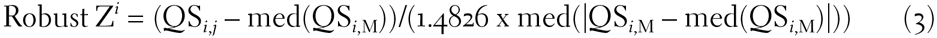

According to the classical and robust Z scores for refining TMR01, TMR04, and TMR07 (Table VII), the best performing forcefields are FF12MC and FF14SBlm; RAPDF/HB_EM_ is better than GBSW; the worst performing forcefield is FF96lm. Both FF12MC and FF14SB refined TMR01, TMR04, and TMR07 substantially better than FF96lm. FF12MC outperforms RAPDF/HB_EM_ and GBSW according to all reported QSs of RAPDF/HB_EM_ and GBSW (sseRMSD, CαRMSD and GDT-TS) listed in Table VI and both classical and robust Z scores listed in Table VII. FF14SB_lm_ also outperforms RAPDF/HB_EM_ and GBSW according to both classical and robust Z scores (Table VII). FF12MC has a robust Z score of 1.33 and a classical Z score of 0.63, while FF14SB_lm_ has both classical and robust scores of 0.04 (Table VII). In terms of refining CASPR models TMR01, TMR04, and TMR07, the present study shows that an improvement of AMBER forcefields has been made from a robust Z score of -0.56 for FF96lm to 0.04 for FF14SBlm and 1.33 for FF12MC over the past two decades. Further, both robust and classical Z scores suggest that FF12MC can generate conformations that cluster around the native conformation of a test protein better than FF14SB_lm_, consistent with the unique abilities of FF12MC to simulate the genuine disorder of C14-C38 in BPTI and its mutant and to sample nonnative states of some miniproteins thus enabling autonomous folding of these miniproteins with folding times close to the experimental values.

**Table VII.**
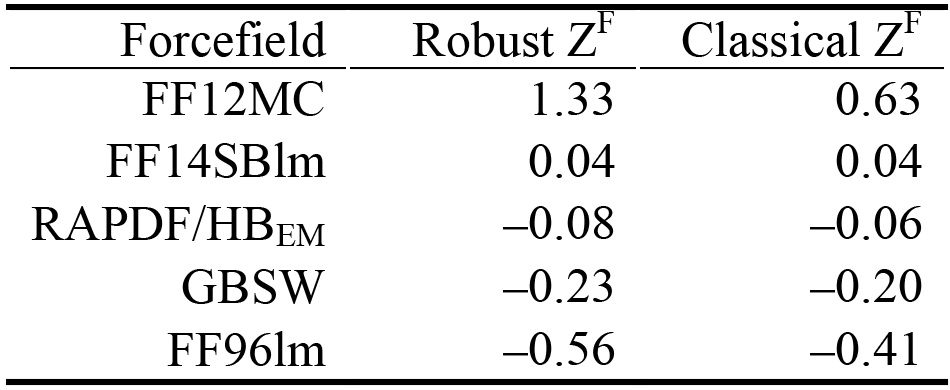
Z Scores for Refining TMR01, TMR04, and TMR07 by Five Forcefields.

### Using FF12MC for protein simulations and known limitations

Confined by current computing speeds, it is challenging to predict the folding kinetics of a miniprotein from MD simulations using the already well-refined, general-purpose forcefield FF14SB^16^ that can fold miniproteins with diverse topologies in MD simulations with implicit solvation.^22^ It is also challenging to use FF14SB to simulate genuine localized disorders of folded globular proteins and to perform simulation-based refinement of comparative models of monomeric globular proteins with large conformational differences from the native conformations. One proposed strategy to take on these challenges is to develop a further-refined specialized forcefield that can sample nonnative states of a miniprotein and localized motions of a folded globular protein without barriers such as certain torsions that are otherwise necessary to achieve agreement between experimental observations and simulations employing implicit solvation. As exemplified above, this type special-purpose forcefield may enable (*i*) capturing the major folding pathways of a miniprotein and thereby correct prediction of the native state conformation and the folding kinetics of the miniprotein, (*ii*) predicting genuine localized disorders of folded globular proteins, and (*iii*) refining comparative models of monomeric globular proteins. The first pursuit of this strategy has culminated in FF12MC.

As a first-generation forcefield specialized for protein simulations with explicit solvation, FF12MC has the following known weakness and limitations. As listed in Table II, FF12MC cannot reproduce main-chain *J*-coupling constants of folded globular proteins as reliably as FF14SB. All bonds involving hydrogen must always be constrained in the NPT MD simulations using FF12MC at Δ*t* = 1.00 fs^smt^ and a temperature of ≤340 K because atomic masses of FF12MC are reduced uniformly by tenfold. FF12MC is not suitable for studying the anomeric effect^153^ or calculation of the entropy of restricted rotation about a single bond^154^ since all torsion potentials involving a nonperipheral *sp*^3^ atom are set to zero. FF12MC should not be used for MD simulations employing PMEMD_CUDA of AMBER 12 or 14 (University of California, San Francisco) without re-compiling PMEMD_CUDA with 0.1008 Da for hydrogen (*viz*., sim.massH = 0.1008 in gputypes.cpp). Preliminary studies showed that FF12MC could fold chignolin from a fully extended backbone conformation to its native conformation in NPT MD simulations performed entirely on a graphics-processing unit (Nvidia GeForce GTX Titan) using PMEMD_CUDA of AMBER 12 with the SPFP or DPDP precision model. When using the SPFP model, there was at least a 6-fold performance improvement of an NPT MD simulation of chignolin performed entirely on an Nvidia GeForce GTX Titan relative to the simulation performed with 16 Intel Xeon E5-2660 core processors (2.20 GHz). However, without an adequate test using the latest PMEMD_CUDA, FF12MC is not suitable for simulations to be performed on graphics-processing units. Instead, FF12MCsm may be experimented on MD simulations using PMEMD_CUDA. Also, no study has been done to determine whether FF12MC can be used for MD simulations at Δ*t* >1.00 fs^smt^ by employing the hydrogen mass repartitioning scheme^53–55^ without compromising the ability of FF12MC to study folding kinetics. Lastly, benchmarking FF12MC against quantum mechanical data of local interatomic interactions and experimental structures of proteins in complex with small molecules is required before the forcefield in its present form can be considered suitable for simulations of a protein in complex with a small molecule.

Nevertheless, FF12MC has the following unique abilities. First, FF12MC can simulate flipping between left-and right-handed configurations for C14-C38 of BPTI and its mutant in solution that was observed in the NMR study of BPTI^61^ and the crystallographic studies of the mutant (Table S12).^66,113^ By contrast, FF14SB locks the C14-C38 bond to the right-handed configuration in solution. Second, FF12MC folds chignolin and CLN025 at 277 K with τ_f_s of 153 and 446 ns^smt^, respectively; whereas the corresponding τ_f_s of FF14SB are 550 and 1012 ns^smt^, respectively (Table IV). These τ_f_s suggest that FF12MC can fold a miniprotein in an NPT MD simulation with folding times that are both statistically 2-4 times shorter than those of FF14SB and closer to the experimental values. Third, the TMR01 refinement by 20 15.8-ns^smt^ of NPT MD simulations at 340 K and Δ*t* = 1.00 fs^smt^ using FF12MC improved the CαRMSD from 6.1 Å to 2.5 Å and GDT-HA from 0.593 to 0.717, whereas the refinement by 20 316-ns^sm^ of NPT MD simulations at 340 K and Δ*t* = 1.00 fs^sm^ using FF14SBlm improved the CαRMSD from 6.1 Å to 3.0 Å and GDT-HA from 0.593 to 0.717 (Fig. S4 and Table S6A and B). These results indicate that FF12MC can improve TMR01 at least 20 times faster than FF14SB when both forcefields were used in low-mass MD simulations under the same conditions. Fourth, it took ~175 days for FF12MC to complete one, unbiased, unrestricted, and 8.85-μs^smt^ classical NPT MD simulation that can fold a 20-residue Trp-cage (TC10b) on a 12-core Apple Mac Pro with Intel Westmere (2.93 GHz). Simultaneously and independently performing 30 distinct and independent simulations of this type led to identification of the two-state kinetics scheme as the major folding pathway of the Trp-cage (Fig. 6). Without any prior knowledge of the hazard function for the nonnative state population of the Trp-cage, the τ_f_ of the miniprotein was predicted to be 1998 ns^smt^ (95% CI: 1396-2860 ns^smt^) from the 30 simulations at 280 K (Table IV). This τ_f_ is consistent with the experimentally determined τ_f_ of 1430 ns^smt^ at 300 K.^71^ By comparison, the folding time of the same Trp-cage sequence reported to date is 14 μs^smt^ that was estimated—also without any prior knowledge that the Trp-cage follows a two-state kinetics scheme — from a pioneering 208- μs^smt^ canonical MD simulation performed on a one-of-a-kind extremely powerful special-purpose supercomputer.^155^ Similarly, the simulations using FF12MC predicted the τ_f_ of the CLN025 to be 174 ns^smt^ (95% CI: 112-270 ns^smt^) at 300 K (Table IV). This is closer to the experimentally estimated value (~100 ns^smt^)^70^ than the reported folding time (600 ns^smt^)^155^ estimated from a 106-μs^smt^ canonical MD simulation of CLN025 on the same special-purpose supercomputer. These results suggest that FF12MC has the ability to sample nonnative states of miniproteins in 20-30 distinct and independent NPT MD simulations.

These results also suggest that one can predict *a priori* whether or not a miniprotein folds according to a two-state kinetics or another scheme at a certain rate without knowing the experimental structure of the miniprotein. As exemplified by the afore-described retrospective predictions of the folding schemes and folding rates of the CLN025 and Trp-cage, the prospective prediction can begin with the use of FF12MC to perform 20-30 distinct and independent NPT MD simulations of the miniprotein to obtain 20-30 sets of instantaneous conformations in time. A cluster analysis of all instantaneous conformations from the 20-30 sets can then be done to define the native conformation of the miniprotein according to the average conformation of the largest conformation cluster. A survival analysis using the 20-30 sets of instantaneous conformations in time and the defined native conformation can then be carried out to determine the folding rate and scheme by examining the hazard function for the nonnative state population of the miniprotein. An increase of the number of distinct and independent simulations may be needed in some cases to avoid a wide 95% CI.

These unique abilities of FF12MC notwithstanding its weakness to reproduce main-chain *J*-coupling constants of folded globular proteins suggest FF12MC may complement FF14SB for kinetic and thermodynamic studies of miniprotein folding and investigations of protein structure and dynamics in areas such as (*i*) estimating the folding rate of a miniprotein using survival analysis of at least 20 simulations, (*ii*) computationally determining whether the folding of the miniprotein follows a two-state kinetics scheme or other schemes by examining the hazard function for the nonnative state population of the miniprotein, (*iii*) simulating genuine localized disorders of folded globular proteins, and (*iv*) refining protein models with large conformational differences from the native conformations.

## Acknowledgments

Yuan-Ping Pang acknowledges the support of this work from the US Defense Advanced Research Projects Agency (DAAD19-01-1-0322), the US Army Medical Research Material Command (W81XWH-04-2-0001), the US Army Research Office (DAAD19-03-1-0318, W911NF-09-1-0095, and W911NF-16-1-0264), the US Department of Defense High Performance Computing Modernization Office, and the Mayo Foundation for Medical Education and Research. The author remains in debt to the late Professor Peter A. Kollman for teaching him the minimalist approach to forcefield development during a one-year sabbatical in the Kollman group 1994-1995. The author is also in debt to the late Professor Shneior Lifson for a stimulating discussion on forcefield development during his visit to the Weizmann Institute of Science, Rehovot, Israel in 1996. The author thanks two anonymous reviewers for their comments and suggestions. The contents of this article are the sole responsibility of the author and do not necessarily represent the official views of the funders.

